# Transgenerational chromosome repair in the asexual bdelloid rotifer *Adineta vaga*

**DOI:** 10.1101/2024.01.25.577190

**Authors:** Antoine Houtain, Alessandro Derzelle, Marc Llirós, Boris Hespeels, Émilien Nicolas, Paul Simion, Julie Virgo, Thomas Lenormand, Bernard Hallet, Karine Van Doninck

## Abstract

Homologous recombination is an essential DNA repair mechanism that promotes chromosome pairing and ensures allele segregation during meiosis in sexual organisms. Here, we explore the dual function of homologous recombination in the bdelloid rotifer *Adineta vaga*, an ancient asexual species known for its remarkable resilience to extreme genotoxic stresses. Genomic analyses reveal that *A. vaga* uses meiotic recombination, both to promote spontaneous crossovers and gene conversion during oogenesis and to repair the genome in response to DNA damage. Our study identifies a novel transgenerational DNA repair mechanism, termed break-induced homologous extension repair (BIHER), which operates on single DNA ends to repair fragmented chromosomes over successive generations. Our findings suggest that meiotic BIHER, coupled with the holocentric structure of chromosomes, represents a key adaptation of life in extreme environments.

## Introduction

Homologous recombination (HR) is a universal mechanism that functions as a high-fidelity DNA repair process of double-strand breaks (DSBs). Defects in HR generally result in cell death or severe dysfunction, including the development of cancer (*1*). In sexual organisms, HR repair of programmed DSBs induced during meiosis is necessary for proper pairing and segregation of homologous chromosomes (*2*). Moreover, meiotic HR generates genetic variability by reassorting alleles along the chromosomes, limiting selective interference between loci and improving the response to selection (*3–5*).

In this study, we investigated the evolutionary and DNA repair functions of HR in the context of an asexual and extremo-tolerant organism, the bdelloid rotifer *Adineta vaga*. These microscopic animals are notorious ancient asexuals, having diversified into hundreds of morphospecies without sex. Additionally, they display a remarkable tolerance to extreme conditions, including complete desiccation and high doses of ionizing radiation (IR), that cause hundreds of DSBs along the chromosomes (*6–9*). The mechanisms that allow their genome to be reassembled are, however, still unknown.

Our recent work has shown that the oogenesis of *A. vaga* is characterized by a modified meiosis I during which homologous chromosomes pair but do not segregate, producing unreduced eggs (*10, 11*). Cytological detection of chromosome pairing was found to coincide with germline DNA repair activation in response to IR treatment, suggesting that HR between homologous chromosomes could be key to maintaining genome integrity (*11*). Furthermore, natural populations of bdelloid rotifers showed indirect genetic signatures of recombination, which supports that HR is involved in their genome evolution (*10, 12–14*).

Here, we surveyed the genome dynamic of *A. vaga* lines in the course of a micro-evolution (ME) experiment spanning 80 to 130 generations, with bottlenecks at each generation following the design of a mutation accumulation experiment. Separate lineages were maintained hydrated or periodically desiccated (Supp. Fig. 1). The implication of HR in genome repair (GR) was further examined by inducing acute DNA damages upon prolonged desiccation or proton IR treatment (Supp. Fig. 2). In both experiments, the genomes of descendants were compared to the ancestor, looking for loss of heterozygosity (LOH) as a signature of HR.

## Results

### Meiotic recombination in the germline

The ancestral laboratory strain of *A. vaga* has a diploid genome structure containing 6 pairs of homologous chromosomes with an average heterozygosity of 1.7% (*10*). In the ME experiment, 11 lines remained hydrated throughout, while another 11 lines experienced 4 cycles of 10-day desiccation (Supp. Fig. 1). All lines were sequenced at the endpoint of the experiment (i.e., after *∼*75 to *∼*135 generations for desiccated and hydrated lines, respectively). Additionally, 14 of these lines were sequenced at the experiment midpoint (after *∼*40 to *∼*70 generations, respectively) (Supplementary Materials). Selection was reduced by collecting a single descendant at each generation. For both hydrated and desiccated lines, the average heterozygosity decreased with the number of generations, with an average loss of 4.23% at endpoint (Fig. 1**a**). However, desiccated samples accumulated loss of heterozygosity (LOH) significantly faster than hydrated samples (Fig. 1**b**), suggesting a possible impact of desiccation on the recombination rate.

**Figure 1:**
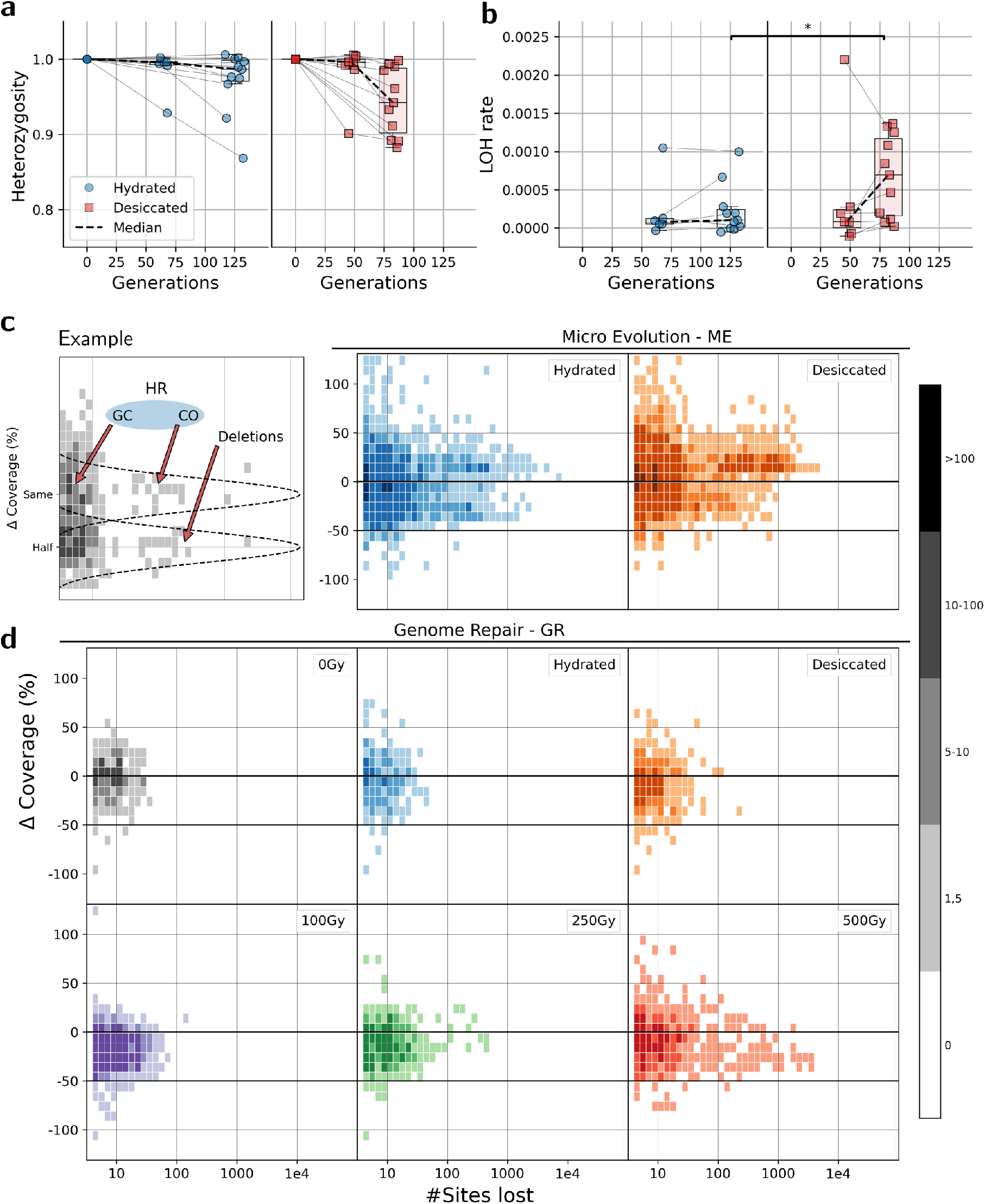
LOH accumulation in ME and GR. **a**: Heterozygosity across generations in both hydrated and desiccated *A. vaga* ME samples. **b**: LOH rate over generations in these ME samples (Mann-Whitney U rank test; p=0.03; CLES=0.22; RBC=0.55). **c-d**: Histograms showing the size distribution of LOH clusters, corresponding to the number of heterozygous sites lost (logged *x−*axis), the frequency (color, *z−*axis), and the Δ% coverage (*y−*axis) for each experimental condition in ME (**c**) and in GR (**d**). In (c), an example illustrating where LOH resulting from deletions and HR would appear on the histogram.

In these 22 ME lines, LOH clusters were detected (Fig. 1**c**). The size of these clusters, determined by the number of contiguous heterozygous sites lost, and their coverage, based on normalized sequencing depth within each cluster, were measured. Clusters with fewer than 5 contiguous sites were ignored, as they could arise from independent LOH events rather than from a genetic event (Supp. Fig. 3). The majority of LOH clusters (85.2%) in ME were smaller than 100 sites. Clusters smaller than 100 sites correspond to LOH tracts that span, on average, less than 5 Kb. These clusters represent only a small fraction of the total LOH (14.2%), whereas clusters larger than 100 consecutive sites (14.8%) account for 85.8% of the total LOH. Additionally, 19.4% of these large clusters (*>*100 sites) correspond to putative deletions (Δ coverage below -20) and make up 11.4% of the total LOH (Fig. 1**c**).

In all samples, allele frequency (AF, Supp. Fig. 4) and coverage are more variable towards the chromosome ends due to a higher content of repetitive DNA (*10*) (as exemplified for chromosome 6 in Supp. Fig. 5). Indeed, sequencing reads tend to misalign between paralogous copies of repetitive sequences, creating a more artefactual heterozygosity signal, correlating with deviations in coverage. To reduce noise, we calculated a LOH score by measuring the AF difference between the ancestor and offspring over a 50 to 100Kb window, corresponding to 1/250 of the chromosome length. A LOH score near 1.0 indicates a LOH event, while a LOH score around 0.0 suggests no detectable change (Supp. Fig. 4**b**). Long LOH tracts, ranging from 50 Kb to over 1 Mb, were identified in 7/11 hydrated lines and in 3/11 desiccated lines (Figs. 2 and 3). Some of these tracts extend to the chromosome ends, while others are found within. For instance, Fig. 2**a** shows a LOH tract of *∼*3 Mb (1) located at the extremity of chromosome 6 in sample D5C3, and another of *∼*115 Kb (2) within an internal region, both detected at the midpoint and endpoint of the experiment. Additionally, smaller LOH tracts (*<*50 Kb) were identified at both sequencing timepoints, as exemplified by tract (3) in Fig. 2**b** (other examples given in Supp. Fig. 6).

**Figure 2:**
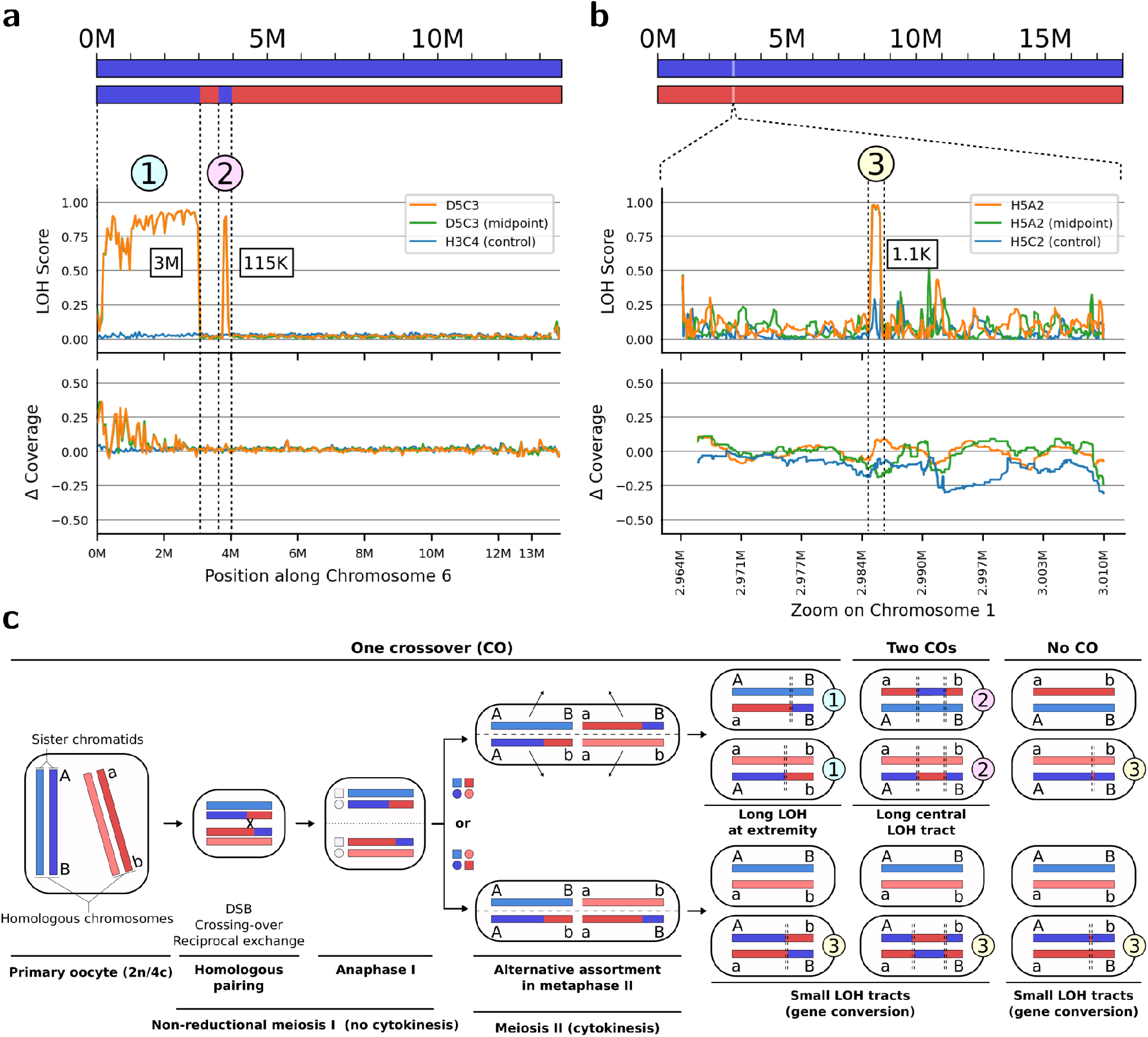
Examples of LOH tracts caused by HR in ME samples. **a**: LOH score and Δ coverage along chromosome 6 of ME sample D5C3 (midpoint and endpoint), illustrating LOH tracts resulting from one CO (example 1) or two COs (example 2). ME sample H3C4 serves as a negative control, with no LOH at the considered positions. **b**: LOH score and Δ coverage along a small portion of chromosome 1 in ME sample H5A2 (midpoint and endpoint), showing an example of a LOH tract likely formed through gene conversion (GC) (example 3). ME sample H5C2 is used as a negative control, with no LOH at the considered position. **c**: Illustration of oogenesis through non-reductional meiosis I, showing expected LOH tracts following homologous recombination with reciprocal exchange or GC. Potential outcomes are given with one, two, or no COs. In some instances, COs followed by segregation can result in both GC and reciprocal exchange (e.g., the case 3 with only one CO), but only the LOH tract due to GC can be detected.

**Figure 3:**
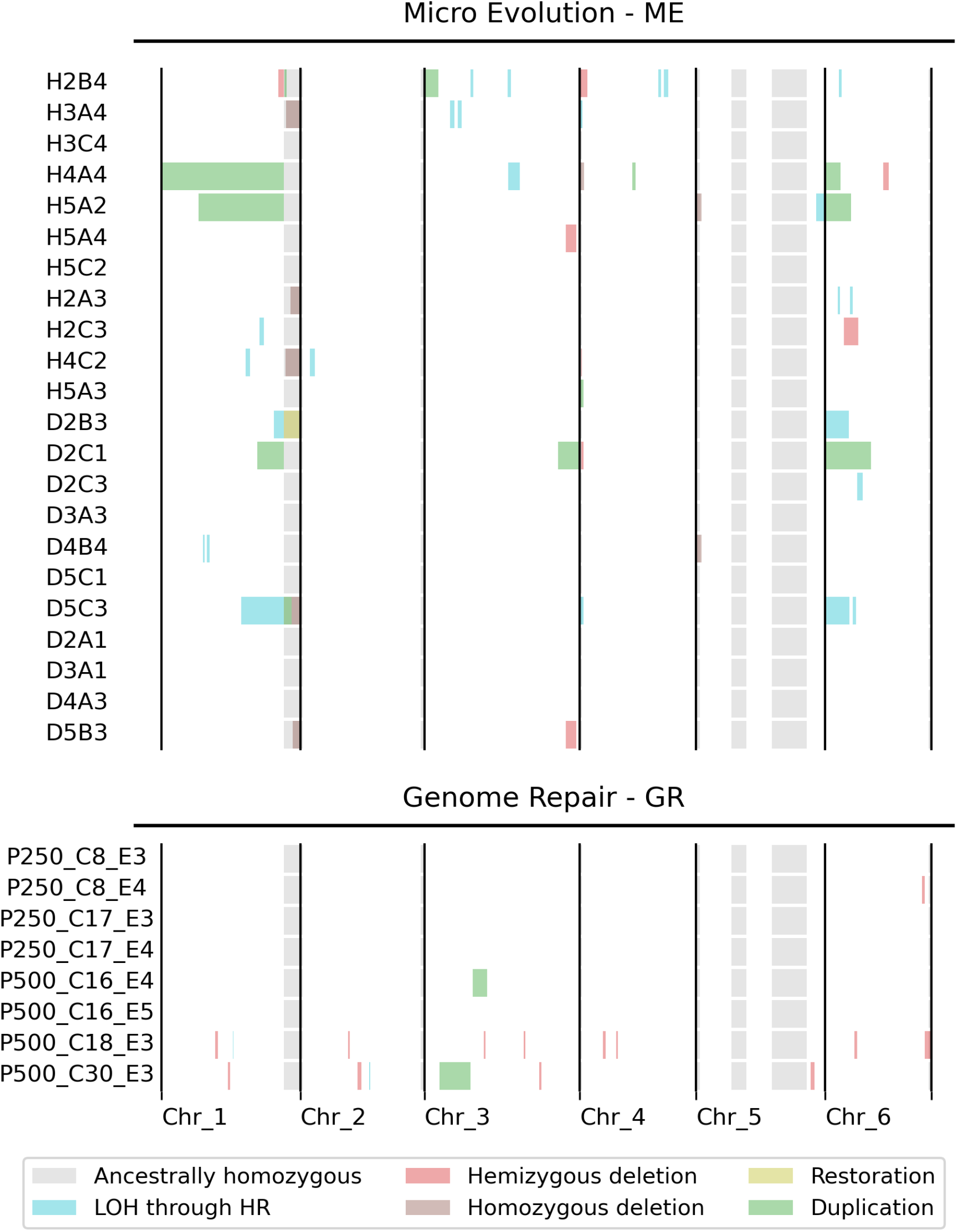
**Summary of the positions of large genomic events** (*>*100 Kb for clarity) observed in ME and GR, categorized per sample and chromosome (based on Supp. Figs. 7 and 14).

These different observations are consistent with HR taking place during the modified meiosis of *A. vaga* when homologous chromosomes assemble into bivalents. Long LOH tracts extending to the chromosome tips are the result of a single crossing-over (CO) (Fig. 2**c**, case 1), while long internal LOH tracts likely arose from a double CO (Fig. 2**c**, case 2). Short LOH tracts (from *∼*100 bp to a few Kb) likely correspond to gene conversion (GC) events inherent to the DNA strand exchange mechanism of HR, even in the absence of CO (Fig. 2**c**, case 3).

All identified long LOH tracts (*>*50 Kb) are present at both the midpoint and endpoint sequencing, except for samples D2B3 and D2C1, where a LOH cluster is found at the midpoint but not at the endpoint (Supp. Fig. 7). In those two cases, HR likely occurred during the culture used for the midpoint sequencing after its separation from the rest of the line.

### Inheritance of deletions and duplications

Besides signatures of reciprocal exchanges between homologous chromosomes, the genomes of ME lines exhibit regions where changes in the LOH score is associated with variations in coverage compared to the ancestor (Supp. Fig. 4**b**; Supp. Table 1). LOH tracts with a decrease in coverage correspond to deletions (Supp. Fig. 8**a-b**), while tracts where coverage increases correspond to duplications (Supp. Fig. 8**c-d**).

Furthermore, deletions are categorized as homozygous when both parental alleles are absent (Supp. Fig. 8**a**) or hemizygous if an allele is absent (Supp. Fig. 8**b**).

Similarly, duplications may represent either the addition of extra copies of specific chromosomal regions (Supp. Fig. 8**c**) or the restoration of diploidy in regions previously affected by ancestral deletions (Supp. Fig. 8**d**).

These homozygous deletions and diploidy restoration likely result from a single CO between homologous chromosomes of different lengths (Supp. Fig. 8**e**). The mechanisms behind hemizygous deletions and other duplications remain less clear, one possibility being unequal crossovers (*15*) (Supp. Fig. 8**f**).

In the 22 sequenced ME lines, we identified 11 large (*>*50 Kb) duplications (including one case of diploidy restoration) and 16 large (*>*50 Kb) deletions, both homozygous and hemizygous, across 13 different *A. vaga* lines (8 hydrated and 5 desiccated). These large genomic events in ME are not randomly distributed but tend to cluster near the chromosome ends (Fig. 3), where transposable element density is higher (Supp. Fig. 9) (*10*). Additionally, samples with recombination signatures accumulated more events than would be expected if they occurred independently (Chi2 = 112.8; p = 5.02e-19) (Fig. 3), suggesting that these recombination signatures are not acquired independently.

Several deletions and duplications were detected at both the midpoint and endpoint of sequencing (Supp. Fig. 7), indicating that these events occurred early in the ME experiment and were transmitted across generations. However, certain regions showed further modifications over time, with changes in Δ coverage present at the endpoint but not at the midpoint. This suggests ongoing recombination within specific chromosomal regions (Supp. Fig. 7). In one instance, chromosome 6 of line H4A4, the changes in AF and coverage are even more complex, indicating a whole genome duplication (WGD) followed by additional duplication of chromosome 1, resulting in aneuploidy, maintained through generations (Supp. Text; Supp. Fig. 10).

### Meiotic HR mediates genome repair in *A. vaga*

In the ME experiment, 10-days desiccation events slightly increased the rate of LOH in *A. vaga* lines, suggesting that desiccation-induced DSBs enhanced HR activity in the germline. To investigate whether HR plays a role in maintaining genome integrity under more severe DNA damaging conditions, *A. vaga* cultures were subjected to longer periods of desiccation (30 days) or irradiated with increasing doses of proton IR (Supp. Fig. 2). Fertility analysis (Supp. Fig. 11) showed that these different treatments only slightly affected the reproductive capacity of *A. vaga*. The most significant effect on growth rate was observed at the highest IR doses of 500 Gy (Supp. Fig. 12), confirming its remarkable resistance to genotoxic stresses (*16*).

The distributions of LOH clusters in non-irradiated (0 Gy) and non-desiccated (hydrated) samples mostly consist of short tracts with fewer than 100 sites, distributed around a Δ coverage of 0 (Fig. 1**d**). As discussed above, these tracts are likely the result of GC. Their presence in untreated samples confirms that recombination between homologous chromosomes occurs spontaneously, independent of external stressors, and the increase in GC signatures in the ME lines compared to the GR samples shows their accumulation over generations (Fig. 1**c-d**). Progeny from the 30-day desiccated cultures had slightly more LOH (4,955 *vs*. 3,169 heterozygous sites lost), with two LOH clusters exceeding 100 contiguous sites (Fig. 1**d**; Supp. Fig. 13), indicating that the DNA damages caused by 1 month of desiccation (estimated at *∼*15–20 DSBs per genome (*16*)) marginally increased the level of HR already present under hydrated conditions. By contrast, IR with 100, 250 and 500 Gy of proton (causing *∼*88, 220, and 440 DSBs per genome, respectively (*16*)) resulted in a marked dose-dependent amplification of the number and size of LOH clusters compared to the non-IR treatment (Fig. 1**d** and Supp. Fig. 13). In particular, 500 Gy IR exposure produced a number of large LOH tracts (*>*100 contiguous sites) distributed among different chromosomes within a same sample (Fig. 3; Supp. Fig. 14). Unlike in ME lines where LOH density increased towards the ends of the chromosomes, IR-induced LOH signatures are more scattered, reflecting the stochasticity of the damage and the level of DNA fragmentation obtained under different conditions (Fig. 3 and Supp. Fig. 9).

### Transgenerational repair of large intra-chromosomal deletions

A large proportion of IR-induced LOH clusters are distributed around low coverage (Δ coverage below -20), indicating that they correspond to deletions (Fig. 1**d**). Specifically, exposure to 100, 250 and 500 Gy IR generated 42.1%, 37.5% and 31.24% deletions, respectively, while in the non irradiated line (0 Gy), deletions represented only 15.85% (Supp. Fig. 13) of all LOH clusters. Most large LOH clusters found in the 500 Gy irradiated lines correspond to hemizygous deletions, in which chromosomal regions up to 400 Kb are lost (Fig. 3; Supp. Figs. 9 and 14).

Deletions may arise from unequal COs as described above for the ME experiment, or be generated by alternative DSB repair mechanisms, such as non-homologous end joining (NHEJ) or single-strand annealing (SSA), which are used to reseal DNA ends while leaving short indels at the repair site (*17*). However, these mechanisms are expected to produce sharp boundaries with an abrupt drop in coverage (coverage change from *∼*1.0 to *∼*0.5) at the breakpoints of the deletion, and a concomitant AF shift (from *∼*0.5 to *∼*1.0) due to the loss of a parental allele (Supp. Fig. 4). This pattern is not observed in the 500 Gy IR deletions. Instead, in all cases, AF progressively increases, stabilizing at *∼*1.0, while coverage gradually decreases towards the centre of the deletion, reaching a minimum of *∼*0.5 to *∼*0.85 depending on the size of the deletion (Fig. 4**a-b**).

**Figure 4:**
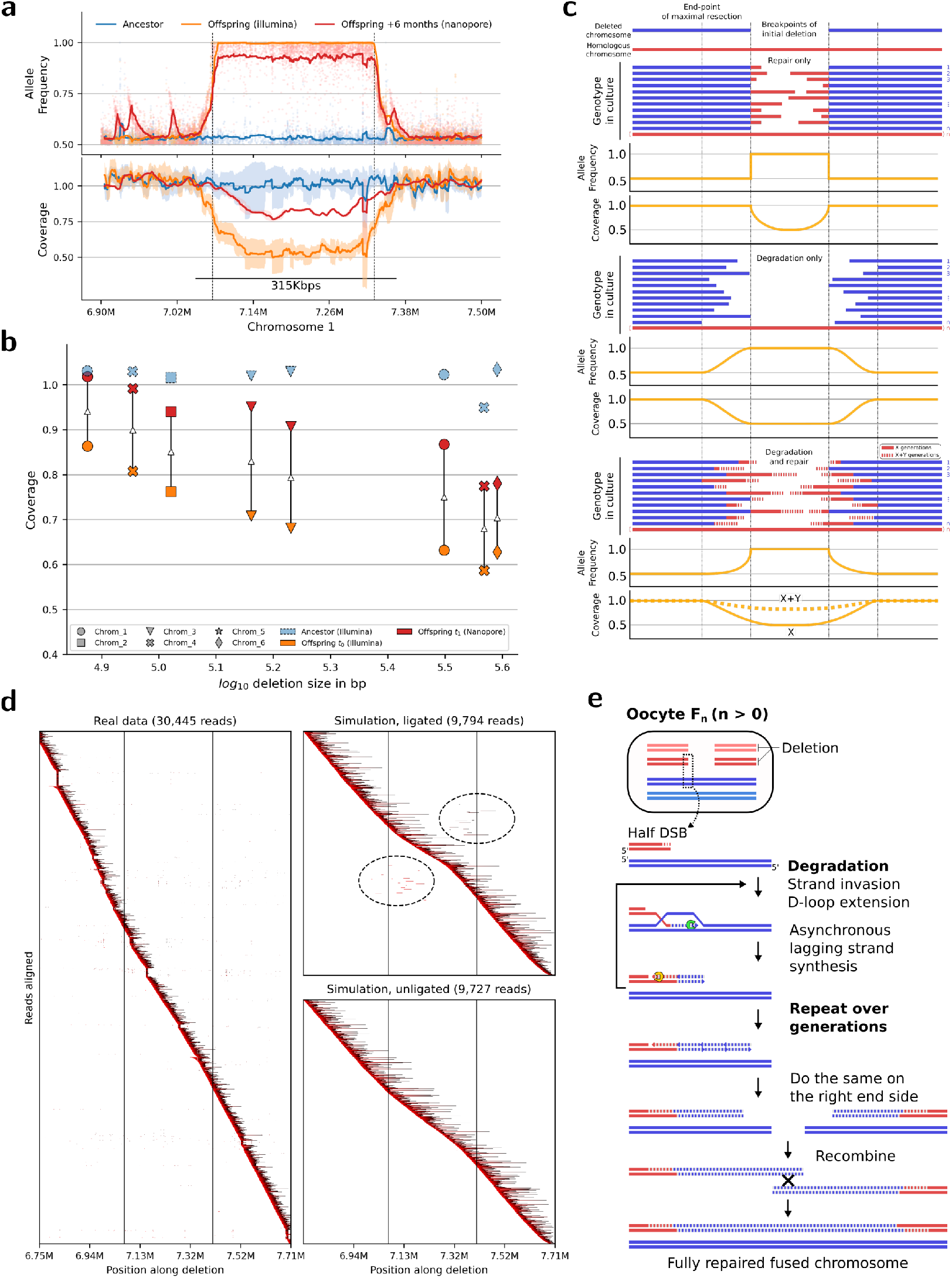
Progressive DNA repair of fragmented chromosomes. **a**: Allele frequency and coverage along a deleted region of chromosome 1 in the GR 500 Gy IR sample P500 C18 E3. **b**: Comparison of expected AF and coverage patterns in a deleted region under scenarios of exclusive repair, exclusive degradation, and a combination of both. **c**: Coverage evolution (*y*-axis) sorted by deletion size (logged *x*-axis) in sample P500 C18 E3, showing two timepoints: Illumina *t*_0_ (blue) and Nanopore *t*_1_ (orange). **d**: Long reads mapped to the reference sequence of the deletion shown in **a**. Alignment color indicates the sequencing read portions that aligned (red: start of the alignment, black: end of the alignment), with the *x*-axis representing the reference sequence position. The vertical lines mark LOH tract boundaries of the deletion. Alignments from the sequencing data (left panel) and both simulated data (right panels, cf. Methods) are shown. **e**: Proposed model of progressive repair of single DNA ends (half-DSB) by BIHER.

This pattern suggests that the descendants of irradiated mothers reduced the deletion to various extents during *∼*1 month of culture used for Illumina sequencing. To test this idea, the 500 Gy IR clone P500 C18 E3, with the highest number of deletions, was grown for an additional six months, and its genome was re-sequenced using Nanopore Technology. The AF profile of the eight P500 C18 E3 deletions remained essentially the same as before (with a plateau at *∼*0.9 instead of *∼*1.0 due to the higher error rate of the sequencing), while the coverage level continued to increase gradually from the edges of the LOH tracts. In the two smallest deletions, coverage reached a value of *∼*1.0, showing a complete recovery of the deleted fragments (Fig. 4**a-b**; Supp. Fig. 15).

These results are consistent with a dynamic process of deletion lengthening and shortening occurring between distant DSBs on a chromosome through generations. Variable DNA degradation from the initial breakpoints first extends the LOH tract, accounting for the AF shift observed at the deletion boundaries. Over time, transgenerational DNA repair outcompetes DNA degradation by gradually replacing the missing fragment using the homologous chromosome as a template (Fig. 4**c**).

After 6 months, the gain in coverage was within the same range for the different deletions (Fig. 4**b**). A diffusion-based mathematical model was fitted to the coverage data and yielded similar estimates for the rates of chromosome degradation and repair (parameters Θ_1_ and Θ_2_) across all deletions (Supp. Fig. 16), supporting the hypothesis that repair was mediated by the same mechanism in all large deletions. This mechanism also accounts for the presence of shorter LOH tracts where AF variability is not accompanied by coverage loss, indicating that in these cases, the initial deletion was fully repaired before Illumina sequencing (Supp. Fig. 17).

### Transmission of broken chromosomes over generations

Intriguingly, alignments of the long Nanopore reads obtained for the partially repaired chromosomal deletions of the irradiated clone P500 C18 E3 show no intermediates in which distal parts of the deleted fragment are joined, suggesting that the boundaries of the deletion were not resealed during the repair process (Fig. 4**d**). Supporting this assumption, *in silico* simulation of 50 resealed intermediates (from Fig. 4**a**) revealed that reads containing re-ligated DNA sequences should have been detected in the alignment if they were indeed present in the actual dataset (Fig. 4**d**; Supp. Fig. 18). This result strongly suggests that IR-induced DNA breakage generated fragmented chromosomes, and that the resulting fragments remained unsealed until fully repaired (Fig. 4**e**).

To confirm the inheritance of broken chromosomes, single-cell eggs from descendants of P500 C18 E3 were arrested in metaphase of first division, and their karyotype was compared to that of the parental strain (Fig. 5). Eggs collected from the control line displayed an average of 12 DAPI-stained bodies, consistent with previous karyotyping and chromosome-level genome assemblies (*10*). In contrast, the average number of DAPI-stained bodies identified in the P500 C18 E3 progeny varied between 12 and 20 (Fig. 5). This result provides conclusive evidence that broken chromosomes can be maintained and transmitted over multiple generations in *A. vaga*. The variable karyotype established for different eggs aligns with the genomic heterogeneity of the P500 C18 E3 line, where large chromosomal deletions were partially or completely repaired over time (Fig. 4**b**; Supp. Fig. 15).

**Figure 5:**
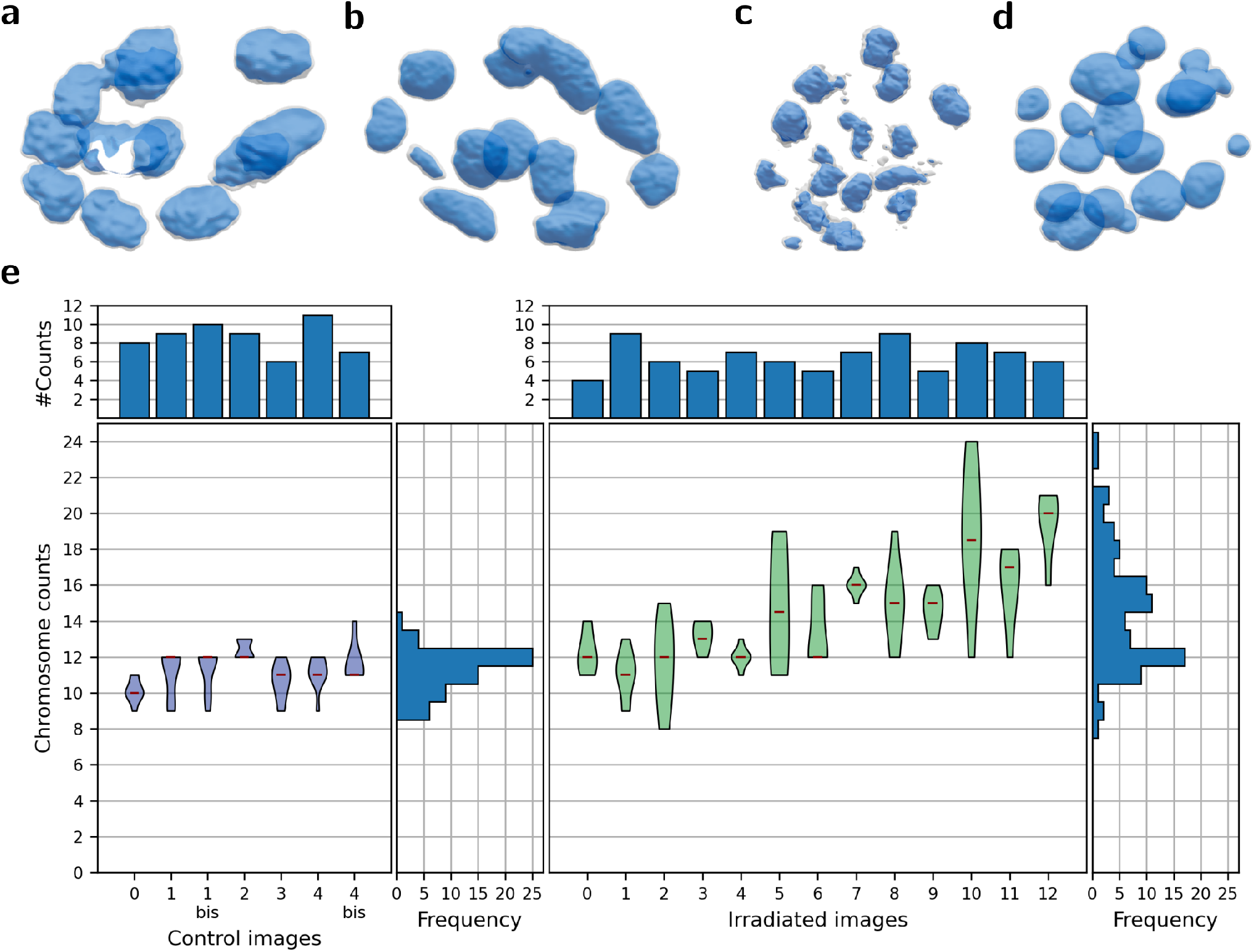
Higher number of chromosome fragments in the descendants of the GR sample P500 C18 E3. **a-b** 3D z-stack microscopy images of a one-cell egg from a non-irradiated control mother, image (a) and (b) correspond to images 1 and 1bis, and 4 and 4bis respectively in panel **e. c-d**: 3D zstack microscopy images of a one-cell egg from a mother of the P500 C18 E3 culture (500 Gy irradiated), image (c) and (d) are represented by images 9 and 11 respectively in panel **e. e**: Independent chromosome counting based on the 3D karyotypes. The top panels display the number of reported counts per image, while the bottom panels show the distribution (violin plots) and medians of counted DAPI-stained bodies per image, along with the frequency of reported counts (histograms) for each experimental condition. Counters were blind to the condition. In control images, some counters were presented with identical images twice (labeled as *bis*) to check consistency. Two out of six counters gave differing counts in only one instance. However, in both cases, the median value of both counts remained identical.

## Discussion

Our previous studies established the diploid nature of *A. vaga*, with evidence for meiotic chromosome pairing, recombination signatures, and delayed DNA repair activity in the germline (*10, 11*). The present study highlights the role of meiotic HR in generating LOH on both a small-scale (GC) and a large-scale (CO).

LOH tracts provide evidence that DSBs, programmed or not, occur frequently in *A. vaga* and induce HR even in the absence of an exogenous source of DNA damage. Desiccation and IR treatment were found to increase the frequency of HR. However, after over *∼*75 generations, LOH accumulated relatively slowly (on average 3.24e-04 per bp per generation in hydrated lines) and remained constrained, (4.23% of heterozygous sites were lost on average and at most 13.2%), despite experimental conditions that minimize selection. The results support the conclusion that HR occurs during non-reductional meiosis I in *A. vaga*, when homologous chromosomes pair, and that segregation of recombinants occurs at meiosis II, when half of the sister chromatids are eliminated through the formation of a polar body (*11*).

Over multiple generations, intra-individual recombination is expected to increase the homozygosity of *A. vaga* lines. While LOH could theoretically unveil deleterious mutations and reduce overall fitness (*18*), experimental evolution studies in highly heterozygous yeasts and theoretical models have shown that LOH can also be beneficial in certain cases, as it can reveal beneficial mutations and reduce the mutational load (*18, 19*). Moreover, intra-individual HR can dissociate linked alleles, reducing selective interference and improving the efficiency of selection, particularly in large populations (*20, 21*).

Intra-individual recombination also allows for new allele combinations by swapping portions of homologous chromosomes. Additionally, HR was found to cause copy-number and structural variations, including deletions and duplications. *A. vaga* lines exhibit high genomic plasticity, tolerating extensive alterations such as aneuploidy and WGD within the timeframe of a few generations (Supp. Fig. 10; cf. Supp. Text). The inheritance of aneuploidy across generations in *A. vaga* is consistent with an oogenesis by non-reductional meiosis I.

One of the most compelling results of our study is the propagation of broken chromosomes arising from IR-induced fragmentation and their progressive reconstruction over multiple generations. This finding points towards a novel transgenerational DNA repair mechanism called break-induced homologous extension repair (BIHER) (Fig. 4**e**). This mechanism is inspired by break-induced replication, which primarily acts to restart stalled replication forks during vegetative growth and telomere maintenance in certain forms of cancer (*22*). We propose that BIHER operates specifically between homologous chromosomes paired during the modified meiosis I of *A. vaga*, coinciding with DNA synthesis signatures that were detected in maturing oocytes following DNA damage induction (*11*). BIHER initiates at one-end DSB (half DSB) resulting from genome fragmentation and subsequent chromosome degradation. The process then proceeds by strand invasion in the template, followed by DNA synthesis, elongating the broken chromosome. Because of limited processivity, elongation stops before reaching the opposite border of the deletion, and the complementary lagging strand is synthesized asynchronously (Fig. 4**e**). This process occurs at both deletion ends, producing extended chromosome fragments which are inherited through mitosis, until they reach meiosis I of the next generation for a new round of BIHER. Over generations, BIHER-mediated elongation outpaces degradation, leading to a reduction in deletion size and a coverage recovery. Eventually, the extended chromosome fragments overlap and fuse back by recombination (Fig. 4**e**).

Stable inheritance of fragmented chromosomes during cell divisions is difficult to reconcile with classical mechanisms of monocentric chromosome segregation in eukaryotes. Typically, acentric chromosome fragments, lacking the centromere, are rapidly lost as they do not attach to the spindle and cannot separate properly in daughter cells (*23*). Instead, we propose that chromosomes in *A. vaga* have a holocentric structure with kinetochores dispersed along their axis, enabling isolated fragments to segregate independently (*24, 25*). This proposition aligns with previous chromosome conformation capture sequencing data, revealing the absence of centromeric signatures in the contact maps (*10, 26*). Holocentric chromosomes have evolved independently in several plant and animal lineages, in certain systems it serves as a possible adaptation to frequent chromosome breakage (*27*). It is therefore tempting to speculate that a BIHER-like mechanism could also work in these organisms as an additional contribution to their tolerance against genotoxic stress.

Regardless of the centromeric structure, unrepaired chromosome fragments expose new DNA ends that can be recognized as DNA damage. Their transmission requires a partial or total inactivation of the DNA damage checkpoint response to allow cell divisions to proceed despite the presence of DNA breaks (*28, 29*). This attenuation has been observed in various systems, including cancer cells (*30*), *Saccharomyces cerevisiae* (*31*) checkpoint mutants or clinical isolates of the pathogenic fungus *Candida glabrata* (*32*). However, in these cases, the suppression of the DNA damage response leads to increased genomic instability, with aberrant chromosome rearrangements such as translocations and chromothrypsis (*33, 34*). Interestingly, no such genomic instability was detected in *A. vaga*, rather triggering DNA repair.

To prevent end-to-end chromosome fusion by end-joining mechanisms, most eukaryotic cells use telomeric protein-DNA complexes composed of shelterin proteins bound to telomeric repeats synthesized by telomerase (*35, 36*). However, new telomeres can also form at ectopic chromosome breakpoints, as observed during programmed DNA elimination in ciliates or nematodes (*25, 37, 38*), or following the induction of DSBs in yeast, mice, and more recently in humans (*39–41*). In *A. vaga*, telomeric repeats were suggested to be added to truncated chromosome ends after the integration of transposable elements like Athena and related retrotransposons (*42*), suggesting that neotelomere formation might protect unrepaired ends resulting from IR-induced DSBs. However, our attempts to detect newly synthesized telomeric repeats at sites corresponding to large deletion repairs were unsuccessful (cf. Supp. Text). If neotelomeres do form in *A. vaga*, they likely have variable positions and are underrepresented in the sequencing datasets, as they may be removed in successive generations through BIHER. Considering the transgenerational BIHER mechanism, it is plausible that chromosomal ends in bdelloid rotifers are maintained and elongated through HR, potentially involving TE elements. In conclusion, the combination of a non-reductional meiosis I, high genomic plasticity, transgenerational repair via BIHER, and efficient DNA repair proteins (*9, 43*) offers key insights into the remarkable genomic resilience of *A. vaga*, and likely bdelloid rotifers as a whole. Our results also support the hypothesis that holocentric chromosomes evolved as an adaptive response to survive increased DNA damage, particularly in extreme environments (*27*). More generally, this research highlights how chromosomal features and reproductive strategies may contribute to genome plasticity, fostering diversification and adaptability. We propose that similar mechanisms may operate in other extremo-tolerant species with holocentric chromosomes, such as tardigrades and nematodes. By maintaining genomic integrity, BIHER likely serves as a critical adaptation that enables extremophiles to thrive in harsh environmental conditions.

## Data and code availability

Haploid genome assembly, gene annotation, and sequencing reads of the ancestor sample are available under the bioproject accession PRJNA680543. The micro-evolution (ME) sequencing reads are available under the bioproject accession: PRJNA929432. The genome repair (GR) sequencing reads are available under the bioproject accession: PRJNA930383.

Scripts and codes used to perform the analysis are available at: https://github.com/AntoineHo/genome_evolution_bdelloids_scripts

## Special thanks

We thank Dr. Matthieu Terwagne for his help and insights into the DNA repair mechanisms. We thank Dr. Aurèle Piazza and Pr. Anabelle Decottignies for their many insightful comments on the manuscript. We also thank Marc Guérineau for his help in experimental design, Jérémy Berthe for the production of desiccated rotifer patches, and Sébastien Meurant and Catherine Demazy for their help with image acquisition on the confocal microscope. The Morphim imaging platform and the Research Unit for Analysis by Nuclear Reactions (LARN) of UNamur are acknowledged for their technical help.

## Supplementary Materials

Materials and Methods

Supplementary Text

Figs. S1 to S20

Table S1 (S2 available as additional electronic content)

References (*44-53*)

