## Supplementary Material w/o Supp Table 2 for "Transgenerational chromosome repair in the asexual bdelloid rotifer *Adineta vaga*"

### Acknowledgments

**Funding** This project received funding from the Horizon 2020 research and innovation program under European Research Council (ERC) grant agreement 725998 (RHEA) to K.V.D., from the Fédération Wallonie-Bruxelles via an "Action de Recherche Concertée (ARC)" grant (vIRGIN) to K.V.D. and B.Ha. Work in the lab of B.Ha. was also supported by a grant from the "fonds Spéciaux de Recherche" (RADyGen) at UCLouvain. B.H. is funded by the BELSPO PRODEX ESA grant ILSRA-2014-0106. A.H. and A.D. are Research Fellows of the Fonds de la Recherche Scientifique—FNRS.

**Data and code availability** Haploid genome assembly, gene annotation, and sequencing reads of the ancestor sample are available under the bioproject accession PRJNA680543. The micro-evolution (ME) sequencing reads are available under the bioproject accession: PRJNA929432. The genome repair (GR) sequencing reads are available under the bioproject accession: PRJNA930383.

Scripts and codes used to perform the analysis are available at:
https://github.com/AntoineHo/genome_evolution_bdelloids_scripts

**Special thanks** We thank Dr. Matthieu Terwagne for his help and insights into the DNA repair mechanisms. We thank Dr. Aurèle Piazza and Pr. Anabelle Decottignies for their many insightful comments on the manuscript. We also thank Marc Guérineau for his help in experimental design, Jérémy Berthe for the production of desiccated rotifer patches, and Sébastien Meurant and Catherine Demazy for their help with image acquisition on the confocal microscope. The Morphim imaging platform and the Research Unit for Analysis by Nuclear Reactions (LARN) of UNamur are acknowledged for their technical help.

### Supplementary Materials

Materials and Methods
Supplementary Text
Figs. S1 to S20
Table S1 (S2 available as additional electronic content)
References *(44-53)*

### Materials and Methods

**Ancestral strain, and cultures** One *Adineta vaga* individual from the AD008 (COI sequence accession number is KM043184) laboratory strain was isolated and cultured until a dense population was established. This population served as the ancestral line for both the ME and GR experiments. All cultures were maintained in SPA® water and were fed with sterile (0.45 $\mu$m pore-size filtering and further autoclaved for 30 min at 121°C) extract of lettuce juice. Subsequent lineages derived from both experiments were cryopreserved at $-80$°C and stored in $1.5$ mL Eppendorf tubes in SPA water®.

**Micro-evolution (ME) experimental setup** The experimental design for ME is outlined in Supp. Fig. 1. Initially, 120 *A. vaga* individuals were randomly selected and isolated from an ancestral culture. These individuals were divided into two batches, each placed across five 12-well plates, with one individual per well. Five plates were assigned to the H (hydrated) condition, remaining continuously hydrated, while the other five plates were allocated to the D (desiccated) condition, undergoing four cycles of 10 days of desiccation followed by re-hydration. Before each desiccation cycle, the cultures were allowed to grow until they reached *ca.* 1000 individuals to increase the survival rate post-desiccation (*16*).

Every fifth and sixth day (for both conditions when D lines were not in desiccation), plates were inspected for eggs, and one hatchling was randomly selected and isolated to begin the next generation. If the isolated hatchling failed to reproduce, the wells from the previous generations were used as backups, with up to five attempts made before declaring extinction. The number of failed attempts was recorded and considered in the final calculation of the number of generations. To prevent overgrowth of contaminants like bacteria or fungi, the culture medium was refreshed periodically throughout the experiment. Lineages were collected and cryo-stored at two key points: the midpoint, upon reaching isolation number 50 for desiccated and 70 for hydrated lines, and the endpoint, when isolation number 88 (desiccated) and 137 (hydrated) were reached. Due to occasional reproduction failures, the actual generation number differed from the isolation number, between 42 and 52, averaging 48 generations for desiccated lines and between 35 and 69, averaging 64 for hydrated lines at the midpoint. Between 75 and 87, averaging 83 for desiccated lines and between 117 and 134, averaging 128 generations for hydrated lines at the endpoint. Extinct lines were not counted in these averages. Desiccation was induced at isolation numbers 20, 42, 62, and 82. Supp. Table 2 summarizes the actual number of generations reached by each lineage at each isolation step.

**Genome repair (GR) experimental setup** The experimental design of GR is summarized in Supp. Fig. 2. In GR, the ancestral *A. vaga* culture was divided into several treatment groups:

- 30H: Maintained in hydrated conditions for 30 days
- 30D: Subjected to desiccation for 30 days
- 0G: Control group, desiccated for 1 day and placed in the irradiation chamber without being irradiated.
- 100G: Desiccated for 1 day, then irradiated with 100 Gy protons
- 250G: Desiccated 1 day, then irradiated with 250 Gy protons
- 500G: Desiccated 1 day, then irradiated with 500 Gy protons

After each experimental treatment, *A. vaga* cultures were re-hydrated and individual rotifers were immediately isolated in multi-well plates. The third, fourth, and fifth eggs laid by the isolated mothers were collected and isolated in individual Petri dishes. The first two eggs were discarded, since they may have started embryonic development and were not at a unicellular oocyte stage during treatment. The isolated eggs were grown into dense cultures (*ca.* 15,000 adults per plate), which were then split into triplicates. The cultures were washed with SPA water, harvested, and the collected rotifer pellets were cryo-stored. The GR sample naming convention was as follows: C## refers to the clone number (*i.e.:* the number of the isolated mother), E## refers to the egg number (*i.e.:* indicating the order in which the eggs were laid).

**Fertility experiment** The fertility and viability of the cultures were assessed in the GR experiment. The experimental design for the fertility assay is summarized in Supp. Fig. 11). A total of 120 females were isolated in 12-well culture plates (Greiner bio-one®) and kept at 19°C in 2.0 mL SPA water for 15 days, with regular feedings of 5$\mu$L of sterile lettuce juice. The number of adult females and laid eggs per well was recorded after on days 7 and 15. Wells with fewer than 2 adults or signs of fungal or bacterial contamination were discarded from the analysis. For wells with over 10 laid eggs, counts were recorded as 10+. Statistical analysis, including two-sided independent t-tests corrected for unequal variance and Glass' $\delta$ statistics (Supp. Fig. 12), was conducted using `SciPy` (*44*).

**Desiccation protocol and irradiation protocol** Desiccation and irradiation of cultures were performed as described in Hespeels et al. (2014) (*16*). Individuals desiccated for 1 day were submitted to 1.7 MeV proton radiation (delivering 25 keV $\mu m^{-1}$).

**Culture collection of GR lines** Cultures of each experimental condition were collected in 1.5 mL Eppendorf tubes with SPA water in quadruplicate, with each tube containing approximately 15,000 individuals. Tubes were centrifuged, and the pellets stored at $-80$°C. The frozen rotifer pellets were later used for genomic DNA extraction.

**Culture and collection of ME lines for sequencing** ME lines selected for sequencing were fetched from the freezer, and cryopreserved rotifers were thawed at $4°C$. Due to the limited number of individuals collected at the midpoint and endpoint of the ME experiment, the thawed rotifers were cultured until the population reached *ca.* 15,000 adults. The cultures were then incubated overnight in 15 mL SPA water with 150 $\mu$L of PenStrep 10,000 U. Cultures were maintained in 75 $cm^2$ cell flasks or in 100 mm Petri-dishes, each containing 15 mL SPA water. For collection, the Petri dishes or cell flasks were vortexed, transferred to 15 mL Falcon tubes and centrifuged at 3,220 g. Supernatant was discarded and pellets from tubes containing the same sample were pooled, refilled to 15 mL with filtered SPA water (0.45 $\mu$m pore-size) and centrifuged again at 3,220 g. Supernatants were discarded, pellets were washed again with 15 mL of filtered SPA water, centrifuged at 3,220 g and transferred to 1.5 mL Eppendorf tubes, adjusted to 500 $\mu$L, and centrifuged at 16,048 g. The final pellets were cryo-stored for subsequent genomic DNA extraction.

**Genomic DNA extraction** Thawed samples stored at $-80$°C were brought to 4°C, and the culture pellets were used for DNA extraction with the QIAGEN Gentra Puregene Tissue Kit. Rotifer pellets, each in a 1.5 mL Eppendorf tube, were centrifuged at 16,048 g for 5 minutes. The supernatant was discarded, and the pellet was kept on ice for 5 minutes. Then, 300 $\mu$L lysis solution was added to each pellet, incubated for 15 minutes, 1.5 $\mu$L proteinase K solution was added, and the pellets were resuspended and incubated overnight at 56°C. The following day, 1.5 $\mu$L RNAse A was added, and the solution was gently mixed and incubated for 30 minutes at 37°C in a shaking dry bath, followed by 2 minutes on ice. Then, 100 $\mu$L of protein precipitation solution was added, and the tubes were vortexed for 20 seconds. The sample was centrifuged at 16,048 g for 4 minutes at 4°C and the supernatant was transferred into a DNA low bind Eppendorf tube containing 300 $\mu$L of isopropanol. The solution was gently mixed by inversion and centrifuged at 16,048 g for 2 minutes at 4°C. The supernatant was discarded, and the pellet was washed with 300 $\mu$L of cold ethanol 70% (stored at $-20$°C), mixed by inversion, and centrifuged again at 16,048 g for 2 minutes at 4°C. After discarding the supernatant, the pellets were air-dried at room temperature for *ca.* 10 minutes, resuspended in 60 $\mu$L of DNA hydration solution, and stored at 4°C until use.

**Illumina sequencing** PCR-free 250-bp paired-end sequencing was performed at the Genomics Core (UZLeuven) on the Illumina HiSeq2000/2500 and NovaSeq platforms, aiming for a sequencing depth of at least 100X per sample.

**Nanopore sequencing** One GR 500 Gy (P500 C18 E3) sample was cultured for an additional 6 months and subsequently sequenced using an Oxford Nanopore PromethION at KeyGene (Netherlands). Genomic DNA provided to KeyGene was treated with a Circulomics SRE kit to remove smaller fragments. For library construction, 2.5 $\mu$g of Ampure purified genomic DNA was used. PromethION 1D Genomic DNA by ligation SQK-LSK109 library prep kit was used to construct a 1D library from the sample and approximately 800 ng was loaded onto a FLO-PRO002 (R9.4.1 pore) version flow cell. The sample was sequenced on two flow cells of a PromethION P24 in two 72 hr runs, as the first flow cell had a relatively low pore count. Basecalling was performed in real-time on the PromethION compute module using the high-accuracy model from package 20.06.18. A total of approximately 123 Gb of ONT data was generated across the two runs. The sequencing reads that passed quality control ($Q \geq 7$) had N50s of 26,140 and 27,035, respectively.

**Quality Control of WGS data** Genomic reads were analysed using FastQC (*45*) and multiQC (*46*). Detailed information on the resulting `.fastq` files is available in Supp. Tables 2. Some samples exhibited adapter sequence contamination at the read extremities, other reads had poor quality metrics at the ends. Both adapter squences and low-quality bases were trimmed before alignment using `fastp`. Additionally, some libraries contained contaminant sequences, detected by the presence of a second %GC peak. Contaminants were mainly bacteria (mostly Enterococcus and *E. coli*) and fungi (mostly *Saccharomycodes ludwigii*) and were retained in the libraries as they would not align to the reference genome.

**Variant calling of Illumina data** The different steps of the variant calling pipeline were performed with a python wrapper pipeline, `runGATK` available at github.com/AntoineHo/runGATK.

Firstly, the Illumina PE reads were pre-processed using `fastp` (*47*) with parameters:

```
--detect_adapter_for_pe
--trim_poly_g
```

Then, reads were aligned to the reference genome assembly (AV20 (*10*)) using `bwa mem` (*48*) with the following parameters:

```
-K 100000000
-R readgroup
```

The read group used depended on the reads library, and was constructed following this pattern:

```
@RG\\tID:{id}\\tSM:{sample}\\tLB:{id}_{sample}\\tPL:{sequencer}
```

Secondly, the `.sam/.bam` files were processed by `sambamba` (*49*) to filter alignments, keeping only those with a mapping quality higher or equal to 20. Then, filtered alignments were sorted and merged for alignments from different libraries of the same sample. Duplicated reads were marked in the merged alignment file. A final `.bam` file per sample was obtained and used for subsequent analyses.

Thirdly, to obtain a single `.g.vcf` file per sample, `GATK4 HaplotypeCaller` was used to call variants with the following parameters.

```
--heterozygosity 0.02
--indel-heterozygosity 0.002
--max-reads-per-alignment-start 0
--output-mode EMIT_ALL_ACTIVE_SITES
--pcr-indel-model NONE
--emit-ref-confidence BP_RESOLUTION
```

Finally, the `.g.vcf` files from all samples (in one experiment), as well as the ancestral population `.g.vcf`, were joint-genotyped using `GATK4 GenotypeGVCFs` with the following parameters:

```
--heterozygosity 0.02
--indel-heterozygosity 0.002
--all-sites
```

Two final `.vcf` files were produced: a raw `.vcf` containing only the short variants detected, and a second `.vcf` file containing all sites information, including short variants and uncalled or homozygous regions.

For the ME experiment, the joint genotyping was performed once with all the samples, and once without the samples from midpoint.

**Genotype change detection** The variant file, containing all reference sites, was annotated to provide a comparative analysis between the ancestor and offspring genotypes using a custom python script. The added information were, 1) the number of samples sharing a common variant (added as a format field SH), 2) the type of variant (added as a format field TP) such as uncalled, biallelic SNP, biallelic indel, multiallelic SNP, etc. 3) a tag indicating how the site changed compared to the ancestor sample (added as a format field GET) such as C for conserved genotype, L for loss of heterozygosity, N for new heterozygous site. A site was considered multi-allelic if more than two alleles accounted for over 10% of the reads aligning at that position.

**Variant calling of Nanopore data** Nanopore reads were aligned on the reference genome using `ngmlr` (*50*). The resulting alignments were sorted using `samtools` (*51*). The coverage was obtained using the `sambamba depth base` (*49*) command and variants were called using `bcftools mpileup` (*51*) and `bcftools call` (cf. Supp. methods for the parameter set used). The final `.vcf` file was analyzed with custom Python scripts.

`bcftools mpileup` was run with the following parameters:

```
--config ont
--annotate "FORMAT/DP,FORMAT/AD"
--ignore-RG
```

`bcftools call` was run with the following parameters:

```
--c -P 0.1 -v
```

**Allele frequency, coverage, LOH tracts and LOH cluster definition** Allele frequency (AF) was defined as the ratio between the number of reads supporting the most common allele at a given genomic position to the total number of reads aligned at that position. AF was measured at all sites where heterozygosity was detected in the ancestral sample. AF should approximate 0.5 at heterozygous positions and reach 1.0 when only one allele is present. Sequencing depth at each position was determined using `sambamba depth base` command (*49*), and both average and median depths per sample were calculated (Supp. Fig. 19). AF and coverage information of heterozygous sites were parsed from the `.vcf` using `pysam`. In subsequent analyses, coverage was defined as the total number of reads aligned to a given genomic position, divided by the average homozygous coverage in central chromosomal regions of the sample.

The average heterozygosity $H$ was calculated by counting the number of heterozygous sites passing the following filters: 0.35 $\leq$ AF $\leq$ 0.65 and 25% $\leq$ coverage $\leq$ 1000% (% coverage is relative to the sample average). The filtered counts, per sample, were then divided by the genome size. Additionally, in the ME experiment only, the LOH rate was computed as $\frac{1-H}{g}$ where $g$ represents the number of generations in the specific ME line under consideration. Average heterozygosity and LOH rates were compared between hydrated and desiccated ME lines at both the midpoint and endpoint of the experiment. Comparisons were also made between endpoint and midpoint values within each condition. Across all pairwise comparisons (e.g., hydrated vs. desiccated, midpoint vs. endpoint), significance was assessed using Student's t-test without corrections if both samples passed a normality test (Shapiro-Wilk test) and had equal variance (Levene's test). Student's t-tests with correction for unequal variance were performed instead if samples had unequal variances. In samples where normality assumptions failed, a Mann-Whitney U (MWU) rank test was performed instead. Effect sizes were reported as Cohen's d for Student's t-tests and as Rank Biserial Correlation for MWU rank tests.

Loss of heterozygosity were detected as clusters of consecutive heterozygous sites in the ancestor that became homozygous in the offspring. The LOH **clusters** were characterized by three metrics: their genomic position, size (counted in number of ancestrally heterozygous sites), and coverage. The $\Delta$ coverage is calculated as follows: $\Delta C = C_{Off} - C_{anc}$ where $C$ is the average coverage of a LOH cluster (example provided in Supp. Fig. 4**a**).

At the chromosome level, LOH cluster analysis fails because they are too frequently interrupted by repetitive content that leads to erroneous heterozygous calls (paralogous mappings). To effectively visualize long LOH **tracts** on a chromosome scale, AF and coverage were averaged within large, non-overlapping windows ($1/250$ of chromosome size) along the chromosomes. The window sizes for each chromosome were 72,587 bp for chromosome 1, 65,099 bp for chromosome 2, 81,419 bp for chromosome 3, 60,899 bp for chromosome 4, 67,722 bp for chromosome 5 and 55,572 bp for chromosome 6. LOH tracts, as well as duplications and deletions, were identified by observing changes in AF, coverage, LOH score (defined as $2 \times |AF_{Off} - AF_{anc}|$) and $\Delta$ coverage. Supp. Table 1 and Supp. Fig. 4**b** summarizes the expected patterns of AF, LOH score, coverage, and $\Delta$ coverage in various scenarios involving LOH, distinct deletion types and duplications.

LOH tracts and clusters were detected using custom Python scripts, with several filters applied to refine the results. The identified LOH clusters were stored in a dataframe that was used to perform all subsequent analyses and representations. Depending on the specific analysis or representation, different filter criteria were applied. A key filter ensured that LOH clusters were unique to individual samples, meaning clusters shared by more than one sample were excluded to reduce false positives. Clusters with fewer than five sites were also filtered out, based on the results of independent LOH clustering (cf. Independent LOH clustering). Additionally, clusters with coverage lower than (<25%) or higher (>1000%), relative to the sample's average coverage, were removed. For chromosomal-scale representation, clusters separated by less than 1 Kb after filtering were merged, preventing artificial splits caused by paralogous mapping in repetitive genomic regions.

**Independent LOH clustering** Some detected clusters of LOH could actually result from independent losses, such as false negatives or variability in allele sequencing depth, rather than from genetic events such as recombination repair or deletions. We modeled independent losses based on the highest observed LOH frequency (being 0.12 in the desiccated ME sample D5C3). This approach helps determine whether LOH clusters of a given size are likely to arise from random independent losses. We modelled the expected LOH clusters using a binomial distribution (Equations 1 and 2). Then, we computed the probability of observing a cluster of any given size (the cluster size is the number of contiguous sites that lost heterozygosity). The results are described in the Supplementary Results section and shown in Supp. Fig. 3.

$$p_{LOH} = N_{LOH}/N_{Het} \tag{1}$$

Where:

- $N_{Het}$ is the number of heterozygous sites in the ancestor sample.
- $N_{LOH}$ is the number of lost heterozygous sites in a given offspring sample.
- $p_{LOH}$ is the probability for a heterozygous site in the ancestor sample to be lost in a given offspring sample.

$$P = (\, n \,)\, N \cdot p^N (1-p)^{n-N} \tag{2}$$

Where:

- $P$ is the probability distribution of cluster lengths given $p$.
- $p$ is the probability of having a LOH event computed by the number of detected LOH divided by ancestral heterozygosity.
- $n$ is the number of trials.
- $N$ is the number of LOH within $n$ trials.

**Simulation of independent recombination events** To model the distribution of events across ME samples at the endpoint, we simulated the random distribution of 53 independent events (successes) across 22 samples in 10,000 iterations using Numpy. The expected number of samples carrying $n$ recombination events was calculated by averaging the result of all simulations. The average number for each $n$ was used as the expected frequencies under the Null hypothesis that events are independent, and compared to observed values of $n$ with a Chi2 test.

**Diffusion model of deletion repair** The deletions appear to be gradually repaired. This repair process was assumed to follow a diffusion-reaction process. Therefore, we modelled the coverage along the chromosomes at two different timepoints (first Illumina sequencing and second Nanopore sequencing) using a differential equation of coverage $f$ at a position $p$ and time $t$ using the function $f(p,t)$ (Equation 3).

$$\frac{\delta f(p,t)}{\delta t} = \theta_1 \frac{\delta^2 f(p,t)}{\delta p^2} + \theta_2 f(p,t)(1 - f(p,t)) \tag{3}$$

$$\begin{array}{lllll} \text{a)} & f(p,0) = 1 & \text{for} & p<p_{min} & \text{or} \quad p>p_{max} \\ \text{b)} & f(p,0) = 1/2 & \text{for} & p>p_{min} & \text{and} \quad p<p_{max} \\ \text{c)} & f(0,t) = f(1,t) = 1 & & & \end{array} \tag{4}$$

$\theta_1$ represents the diffusion coefficient associated with the repair process and $\theta_2$ the rate of repair, $p_{min}$ and $p_{max}$ are the initial boundary positions of the deletions along the chromosome. The parameters $\theta_1$ and $\theta_2$ were adjusted by least square. The coverage data were smoothed using a moving average with a window width of 50 positions. The coverage data was slightly shifted up or down, so that their average value was near one at both extremities, as assumed in Eq. 4b. This shift was calculated as one minus the average frequency observed in the windows defined from $[0, p_{min}/2]$ to $[1/2 + p_{max}/2, 1]$.

**Deletion and Nanopore reads simulations** `samtools faidx` was used to extract the reference sequence corresponding to a large deletion observed in sample P500 C18 E3 on chromosome 1. Nanopore long reads were aligned using `minimap2` with splice-aware settings (`-ax splice -G400k`). To analyze the error profile of the real Nanopore dataset, `nanosim` (*52*) was used and then two new sets of long reads were simulated based on new reference sequences. The simulated reference sequences were generated by introducing 50 pairs of variable breakpoints around the actual deletion sites to account for observed variability in deletion size. Two sets of long reads were produced per deletion: in one set the deletions were ligated *in silico* (set 1), in the other set, the deletions were left unligated (set 2). In addition, both sets also contained 50 undeleted reference sequences to simulate the presence of a non-deleted allele. The long reads of both sets were aligned with `minimap2` using the same splice-aware settings. A custom Python script was used to plot the alignments obtained with the real dataset and both simulated sets (unligated vs. ligated deletions). A summary of this methodology is presented in Supp. Fig. 18.

**Karyotypes** A clonal culture was initiated using a rotifer back-up from the GR sample P500 C18 E3 and maintained in a 75 $cm^2$ cell flask until the population reached *ca.* 40,000 individuals. A clean laboratory strain culture was similarly maintained and used as a control. The individuals were then concentrated in a smaller volume (10 $cm^2$ Petri dish) and cultured for two days at 25°C in nutrient-rich water supplemented with 10 $\mu$g/mL demecolcine to stop egg development during the first metaphase. After two days, eggs were collected in a 1.5 mL Eppendorf tube and centrifuged at 7,379 g. The supernatant was discarded, and 500 $\mu$L of SPA water (put at pH 10 with NaOH) was added. Then, 500 $\mu$L of a diluted bleach solution (14 mL SPA water with 1 mL bleach) was added. The tubes were vortexed, centrifuged again at 7,379 g for 2 minutes, rinsed with 1 mL of SPA water, and the supernatant was removed. These washing steps were repeated three times. The egg pellet was resuspended in 1 mL of a hypotonic solution (0.05 M sucrose in distilled water) and transferred to a 15 mL Falcon tube. While the tube was vortexed at high speed, 4 mL of the hypotonic solution was slowly added drop by drop. The eggs were incubated overnight on an agitator. The tubes were then centrifuged for 10 minutes at 3,220 g and the supernatant was removed. Next, 5 mL of a fixative solution (3 parts methanol to 1 part acetic acid) was slowly added drop by drop while the tubes were vortexed at high speed. The tubes were left on ice for 1 hour and then centrifuged at 3,220 g for 10 minutes. The supernatant was discarded and 5 mL of the fixative solution was added drop by drop while the tubes were vortexed at high speed. The tubes were left on ice for 20 minutes and centrifuged at 3,220 g for 10 minutes. The supernatant was discarded. *ca.* 15 $\mu$L of the fixed eggs were dropped from about 15 cm onto a microscope slide. The drops were air-dried and, for some slides, gently squashed by pressing on the coverslip. The samples were then stained with DAPI (1 $\mu$g/mL in PBS) and washed twice with PBS. The slides were mounted with fluoromount-G, allowed to solidify for at least one hour at 37°C) and stored at 4°C until observation. Z-stack images of one-cell-stage eggs were then acquired using a Zeiss LSM Airyscan confocal microscope. Images were taken only when chromosomes were clearly visible and when at least 10 chromosomes were initially counted. The Z-stack images were thresholded and converted to 3D models using a custom Python script. These 3D images were uploaded to a public website, where users could input chromosome counts. To maintain objectivity, the experimental condition was masked to the users, and some images were duplicated to check consistency in counting. Final counts were aggregated, noisy images with high variability in counts were discarded from the analysis.

### Supplementary Text

**Coverage evolution over culture time indicates similar deletion repair kinetics across deletions of P500 C18 E3** The coverage dynamics resulting from the degradation and repair of deletions were modeled using data from the irradiated sample P500 C18 E3. A diffusion equation (Supp. Methods) was applied to model the coverage curve, given a degradation rate ($\theta_1$) and a repair rate ($\theta_2$). Best-fit estimations of these rates were obtained for each deletion, and were used to produce predictions of the coverage curve shape (Supp. Fig. 16). The fitted parameter values were consistent across deletions, indicating that the repair and degradation balance is comparable across all deletions. As expected, the repair rate exceeded the degradation rate, suggesting a common repair mechanism across all deletions. Interestingly, some deletions present asymmetric boundaries and exhibited the poorest fit to the model, potentially indicating that the rates of end-resection and repair differ across the length of the chromosomes. Coverage is also affected by local sequence characteristics, e.g., paralogous sequences such as TEs create higher local coverage variance by preventing confident reads alignment. As a consequence, the chromosomal location of a deletion impacts the best-fit estimates, as more TEs are found at the chromosome extremities. When the difference of coverage was too important between both sides of a deletion (typically with lower coverage near the chromosome ends), the data was adjusted before fitting (see Supp. Methods).

**Irradiation impacts individual fertility** To assess the impact of DNA damage on reproduction, a fertility assay was conducted on individuals from cultures that underwent the treatment in the GR experiment. The number of offspring produced by treated mothers was counted after one and two weeks of culture. Both desiccation and IR significantly affected fertility compared to untreated controls (Supp. Fig. 12**a-b**). The growth rate between the second week and first week was used to correct for age variability when the mothers were isolated (Supp. Fig. 12**c**). This correction reduced the effect size and significance for most conditions. However, a significant decrease and strong effect size remained for 500 Gy IR samples ($p = 4.6e^{-05}$ independent t-test corrected for unequal variance, Glass' $\delta \sim 0.6$). Moreover, fewer cultures could be maintained from eggs isolated from 500 Gy IR-treated mothers (6 mothers, 10 eggs) compared to the other experimental conditions (with an average of 11.4 mothers and 27.0 eggs across conditions except for 500 Gy).

**Spontaneous deletions and duplications are transmitted over generations** Deletions and duplications were detected in the micro-evolution experiment as variations in ($\Delta$) coverage that correlate with shifts in allele frequency and LOH score. Deletions are classified as homozygous when both parental alleles are absent (Supp. Fig. 8**a**) or hemizygous if only one allele is missing (Supp. Fig. 8**b**; Expected patterns in Supp. Fig. 4). Among the large deletions (>50 Kb) observed across the ME lines (17), most are located at chromosome ends (12), with 8 being homozygous and a few terminal deletions being hemizygous. The remaining deletions (5) are found in the middle of chromosomes and are hemizygous (Fig. 3) (summary based on Supp. Fig. 7).

An increase in coverage indicates the presence of additional copies of chromosome segments (Supp. Fig. 8**c**). In one example in the ME lines, a coverage increase reflects a restoration of diploidy caused by the duplication of the longer homolog of chromosome 1, where the ancestral homologs are of unequal length (Supp. Fig. 8**d**). The other duplications (13 in total) represent extra copies of distinct genomic regions (Fig. 3; Supp. Fig. 7).

Restorations (1/13) and homozygous deletions (8/17) likely result from a single CO between homologous chromosomes of different sizes. Whether a deletion or a restoration occurs, depends on how the chromatids segregate (Supp. Fig. 8**e**). The underlying mechanism for the remaining deletions and duplications is unclear, but unequal crossovers may play a role (*15*) (Supp. Fig. 8**f**). However, some deletions have patterns of AF and coverage inconsistent with unequal CO. For example, one large deletion has a

LOH tract that extends beyond the coverage drop, suggesting partial repair (Supp. Fig. 20**a-b**). Other deletions display a progressive drop of coverage over dozens of Kb (Supp. Fig. 20**c-d**) where a breakpoint would be expected (e.g., Supp. Fig. 20**e-f**). In one particular case, a deletion had a progressive coverage decrease on one side and a breakpoint on the other side (Supp. Fig. 20**g**).

Large duplications spanning several Mb were identified in four ME lines: H2B4, H4A4, H5A2 and D2C1 (Fig. 3), consistently at chromosome ends. For instance, in H4A4, chromosome 1 appears to be fully duplicated, while another duplication of approximately 1.8 Mb is present at the end of chromosome 6. These duplications in H4A4 were detected at both the 61st and 118th generations (Supp. Figs. 7; 10**a-b**), indicating their early appearance in the ME experiment and stable transmission across generations. By the 118th generation (endpoint), chromosome 1 exhibits further duplications in one region, with a higher $\Delta$ coverage at the endpoint compared to the midpoint (Supp. Fig. 10**a**). Similarly, the duplication on chromosome 6 has expanded by roughly 500 Kb at endpoint compared to the midpoint, suggesting a progressive accumulation of duplications over generations (Supp. Fig. 10**b**).

In most cases, duplications and deletions are likely caused by unequal crossovers between repetitive sequences. However, the mapping of short reads prevents determining whether the duplications are actually in tandem or exist as independent fragments. In one particular case, shifts in coverage and AF (Supp. Fig. 10**c**) correspond to a ratio of 5:4 copies, resulting in a coverage level of 1.25 and an AF of 0.6 (tetraploid), rather than the expected values for diploid (0.5) or triploid (0.66). A single duplication of chromosome 1 (triploidy) cannot explain these observations. Furthermore, the observed AF shifts do not span the entire chromosome. This result suggests that line H4A4 likely underwent a WGD, followed by a duplication of the shorter homolog of chromosome 1.

### Supplementary Figures

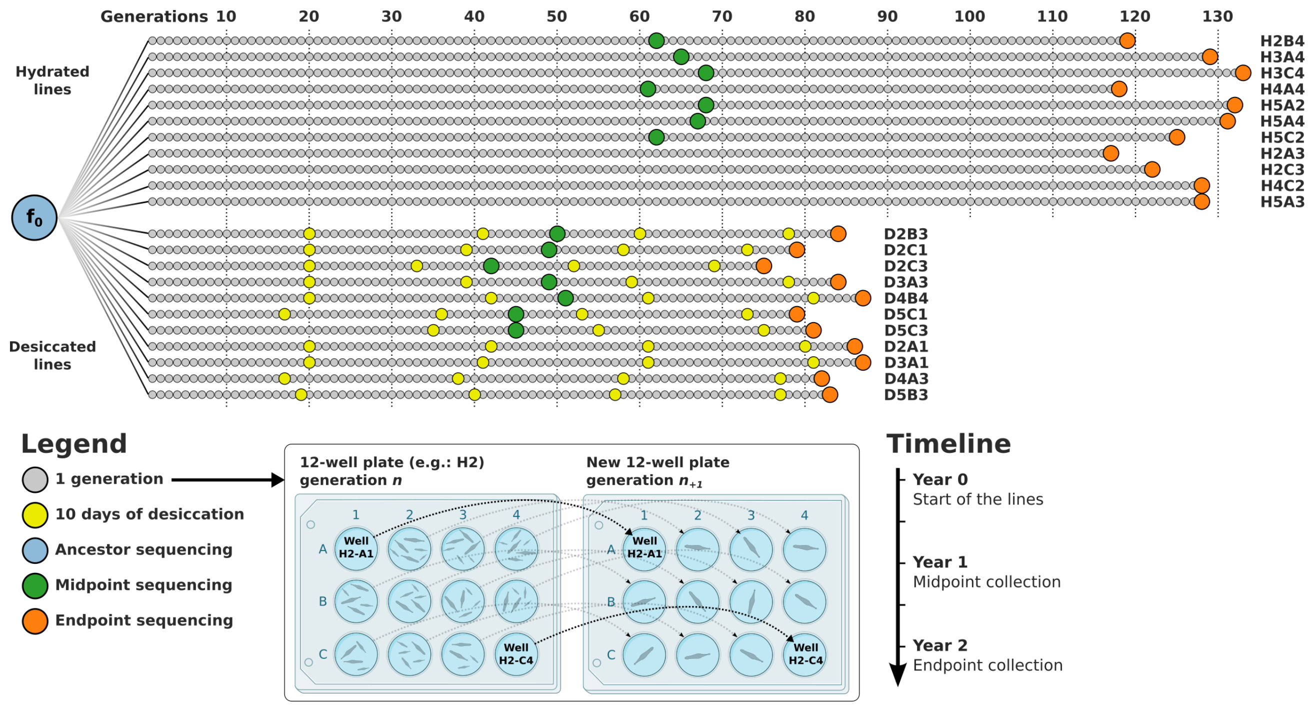


Supplementary Figure 1: Summary of the experimental design of the micro-evolution (ME) experiment. Starting from the same ancestral culture ($f_0$, blue circle), 22 *A. vaga* lineages were experiencing bottlenecks at each generation, following the design of a mutation accumulation experiment (*53*). Those 22 lines were sequenced using Illumina 2x250 bp PE after 75 to 135 generations at the experiment's "endpoint" (orange circles). In addition, 14 lines were sequenced after one year of the experiment, corresponding to 40 to 70 generations at "midpoint" (green circles). Two types of treatments were applied, **H**(ydrated) lines were kept in permanent hydrated conditions. **D**(esiccated) lines underwent four desiccation events of 10 days (yellow circles) roughly every 20 generations.

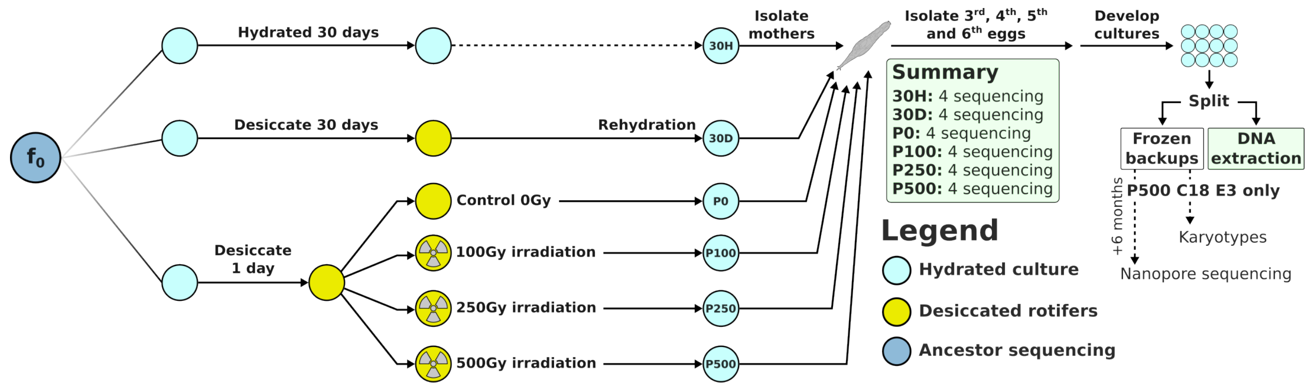


Supplementary Figure 2: Summary of the experimental design of the genome repair (GR) experiment. All cultures originated from the same ancestral laboratory strain ($f_0$, blue circle). Cultures were divided into experimental treatments: one month under hydrated conditions served as the control (30H), one month under desiccated conditions (30D), and exposure to proton irradiation at doses of 0 (control P0), 100 (P100), 250 (P250) and 500 (P500) Gy. Individual mothers were isolated from the treated cultures, and the third, fourth, fifth eggs laid by those mothers were collected individually to initiate new cultures (one-egg bottleneck). Four cultures from each treatment were sequenced using Illumina 2x250 bp PE. One of the four samples sequenced from the P500 series (P500 C18 E3) was used to generate microscopy images of single-celled egg karyotypes and was later re-sequenced using Nanopore long reads technology after 6 months of culture.

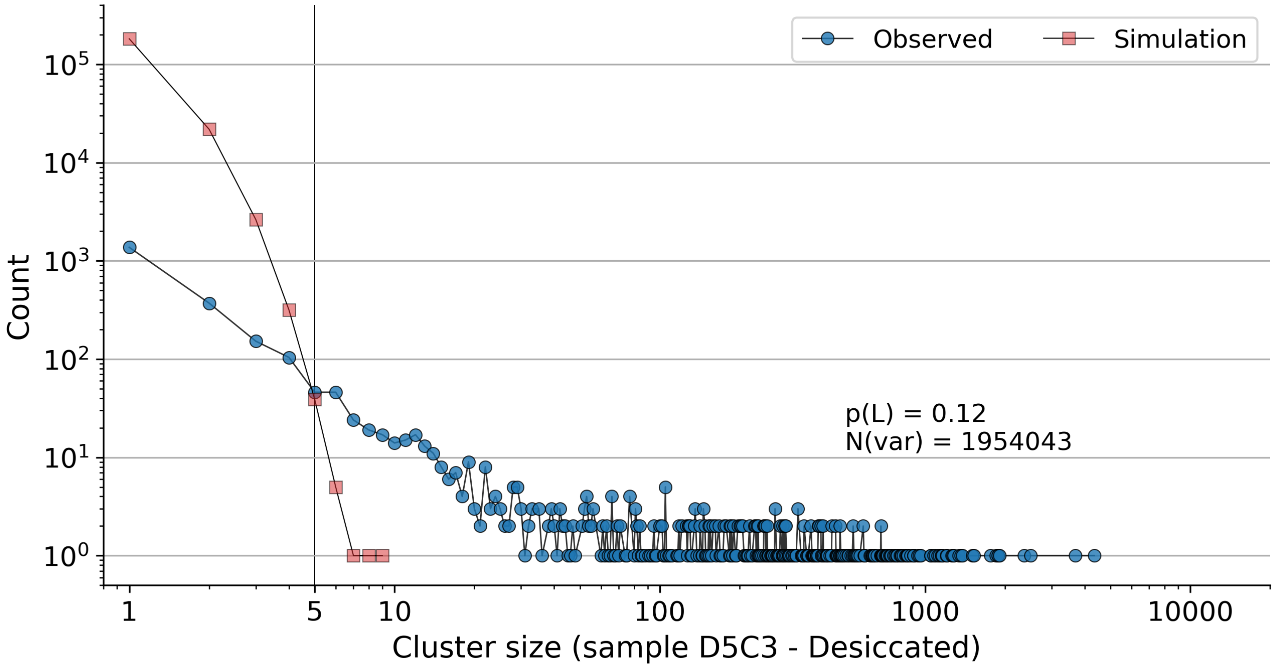


Supplementary Figure 3: The number of LOH clusters of a given size (measured by the number of sites lost) was compared between observed data and simulated data, using a model of independent LOH events. The simulation was based on a binomial distribution with $p_{LOH} = N_{Hom}/N_{Het}$.

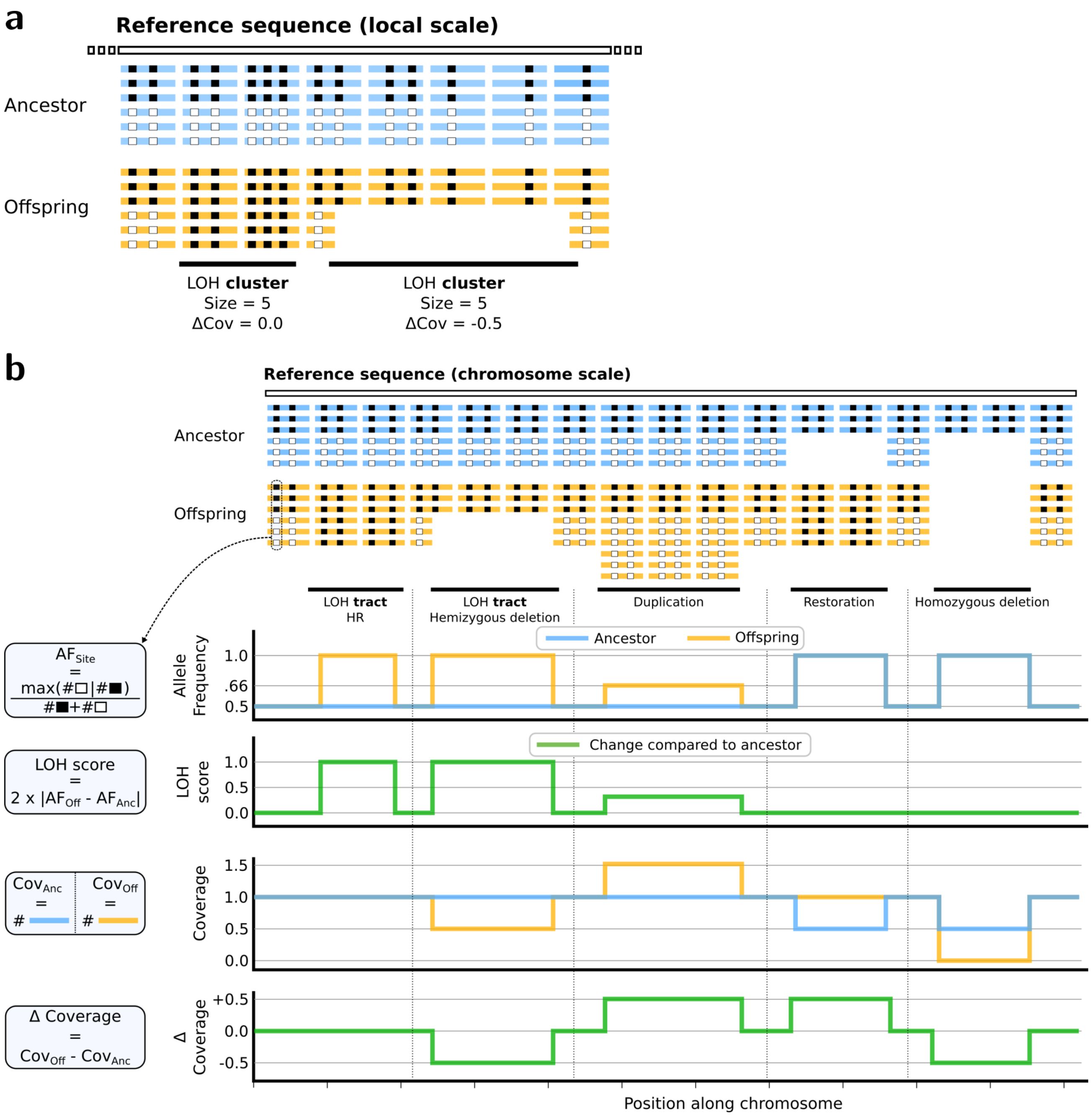


Supplementary Figure 4: **a**: Signatures of LOH clusters resulting from homologous recombination (HR) or deletions. **b**: Detection of LOH tracts, hemizygous and homozygous deletions, duplications, and restorations based on patterns of allele frequency (AF), coverage, LOH score and $\Delta_{Coverage}$.

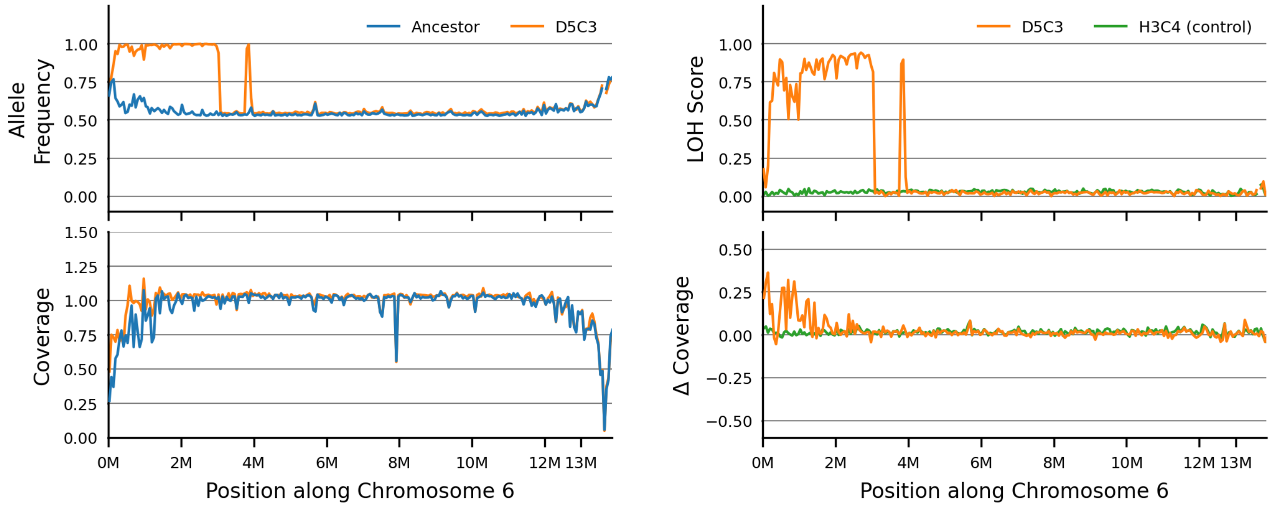


Supplementary Figure 5: Left panel, allele frequency (AF, upper graph) and coverage (lower graph) along chromosome 6 in the ancestral strain and ME sample D5C3. Right panel, LOH score (upper graph) and $\Delta$ coverage (lower graph) along chromosome 6 in ME samples D5C3 and H3C4 (this latter sample is used as a control with no LOH on this chromosome).

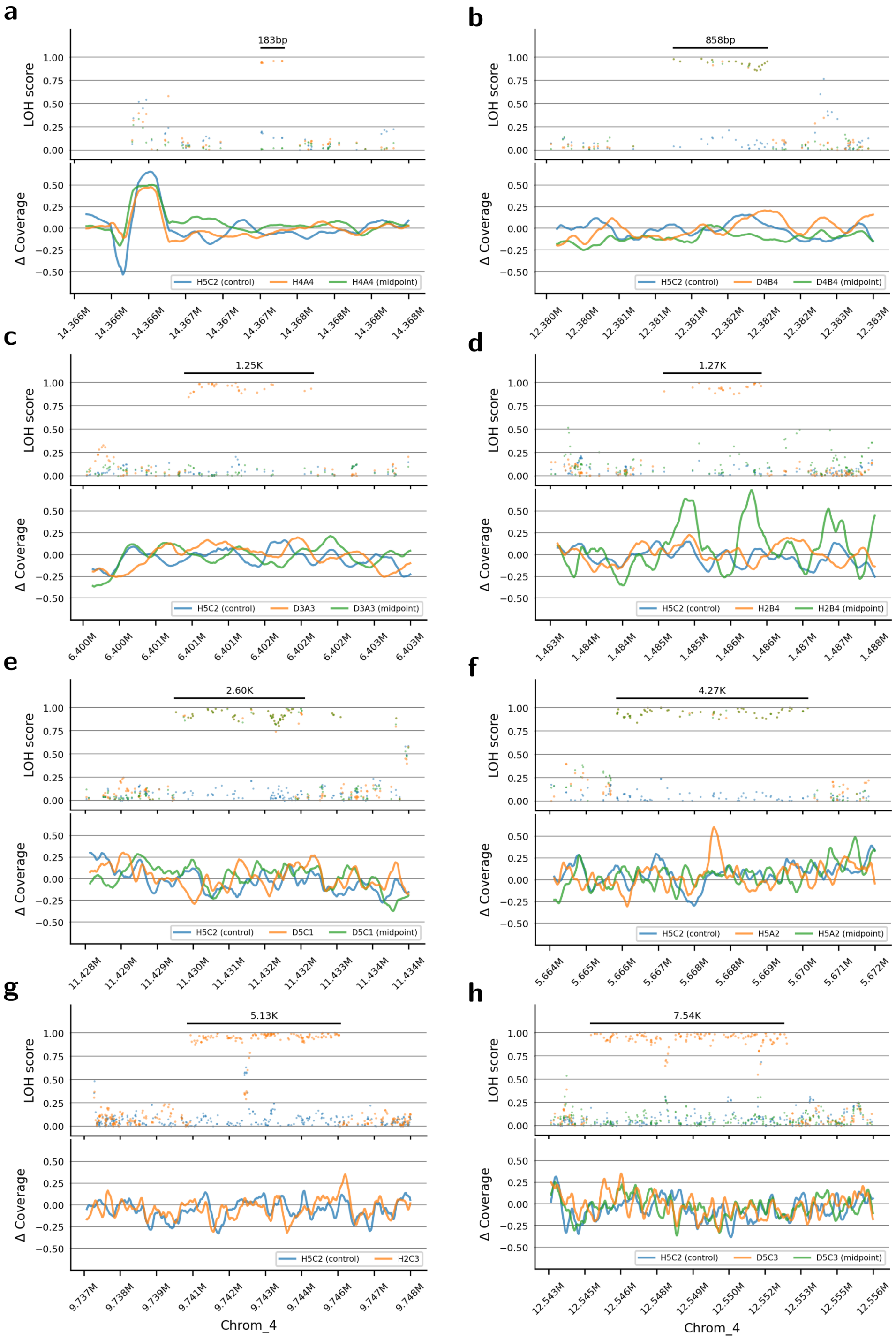

Supplementary Figure 6: **a-h**: LOH score and Δ coverage for representative examples of short LOH tracts (<50 Kb) identified in ME samples.

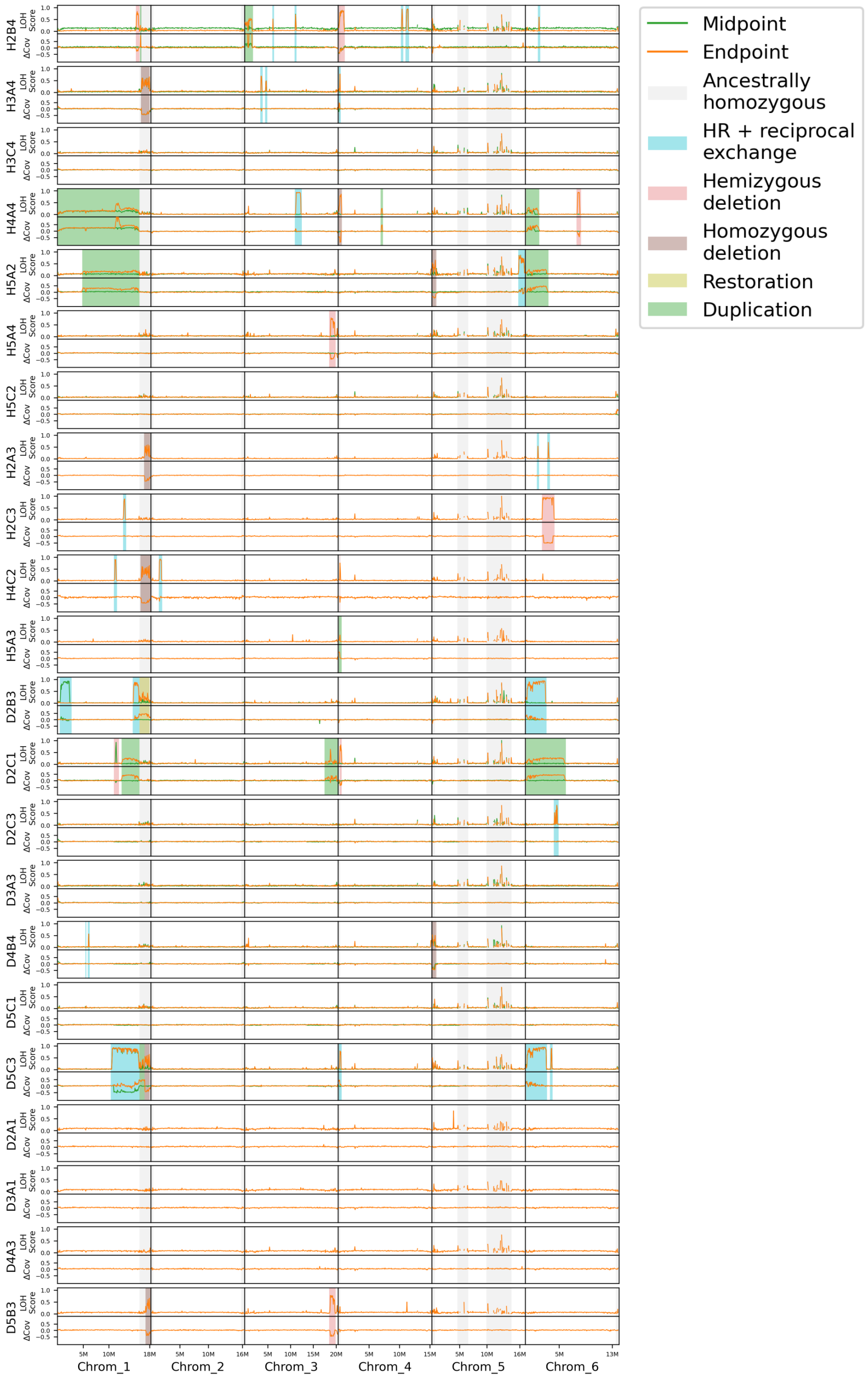


Supplementary Figure 7: LOH score and Δ coverage across all chromosomes in ME samples with large events (>50 Kb) highlighted in color: blue for HR and reciprocal exchange, light red for hemizygous deletions, dark red for homozygous deletions, yellow for restorations, and green for duplications. Grey areas represent ancestrally homozygous regions. Values are averaged over 250 non-overlapping windows along each chromosome, with window sizes comprised between 55.6 Kb (Chromosome 6) to 81.4 Kb (Chromosome 3), depending on chromosome length.

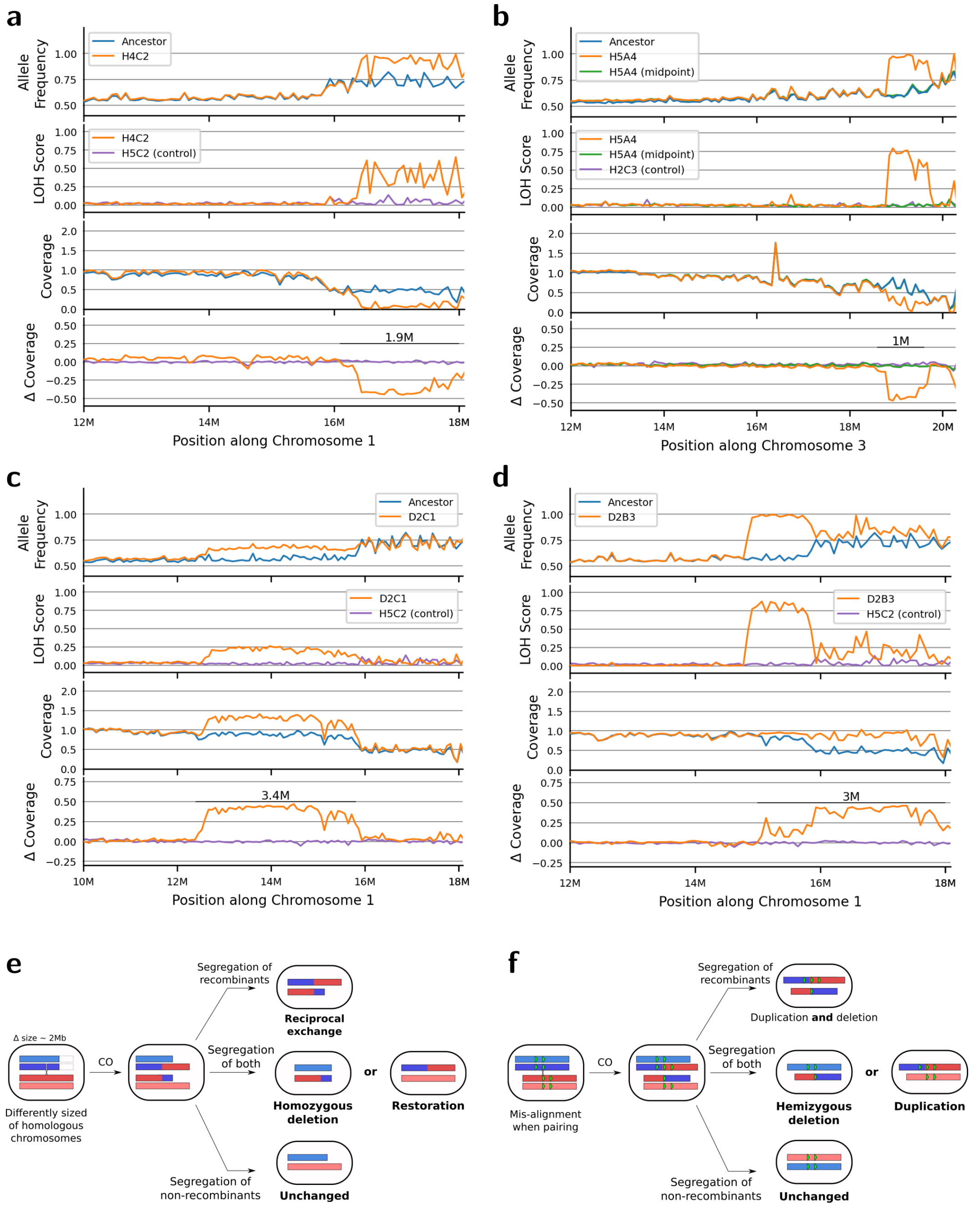


Supplementary Figure 8: **Large deletions and duplications observed after multiple generations in the ME experiment. a-d**: Allele frequency, LOH score, Coverage, and Δ coverage across non-overlapping windows at chromosome ends. **a**: Homozygous deletion on chromosome 1 in sample H4C2 (window = 72.6 Kb). **b**: Hemizygous deletion on chromosome 3 in sample H5A4 (window = 72.6 Kb). Only 4 hemizygous deletions were detected at chromosome ends, and due to alignment noise, they are difficult to distinguish from homozygous deletions. Coverage retention was used to confirm these hemizygous deletions, which retained coverage, whereas homozygous deletions showed complete coverage loss. **c**: Duplication of a region on chromosome 1 in sample D2C1 (window = 81.4 Kb). **d**: Restoration of an ancestral deletion on chromosome 1 in sample D2B3 (window = 72.6 Kb). **e**: Recombination between homologous chromosomes of unequal sizes (e.g., chromosome 1 of *A. vaga*) can produce homozygous deletions or restorations of coverage at chromosome extremities, depending on chromatid segregation. **f**: Recombination between mis-aligned chromosomes (i.e.: at repetitive regions) can produce segmental duplications or deletions.

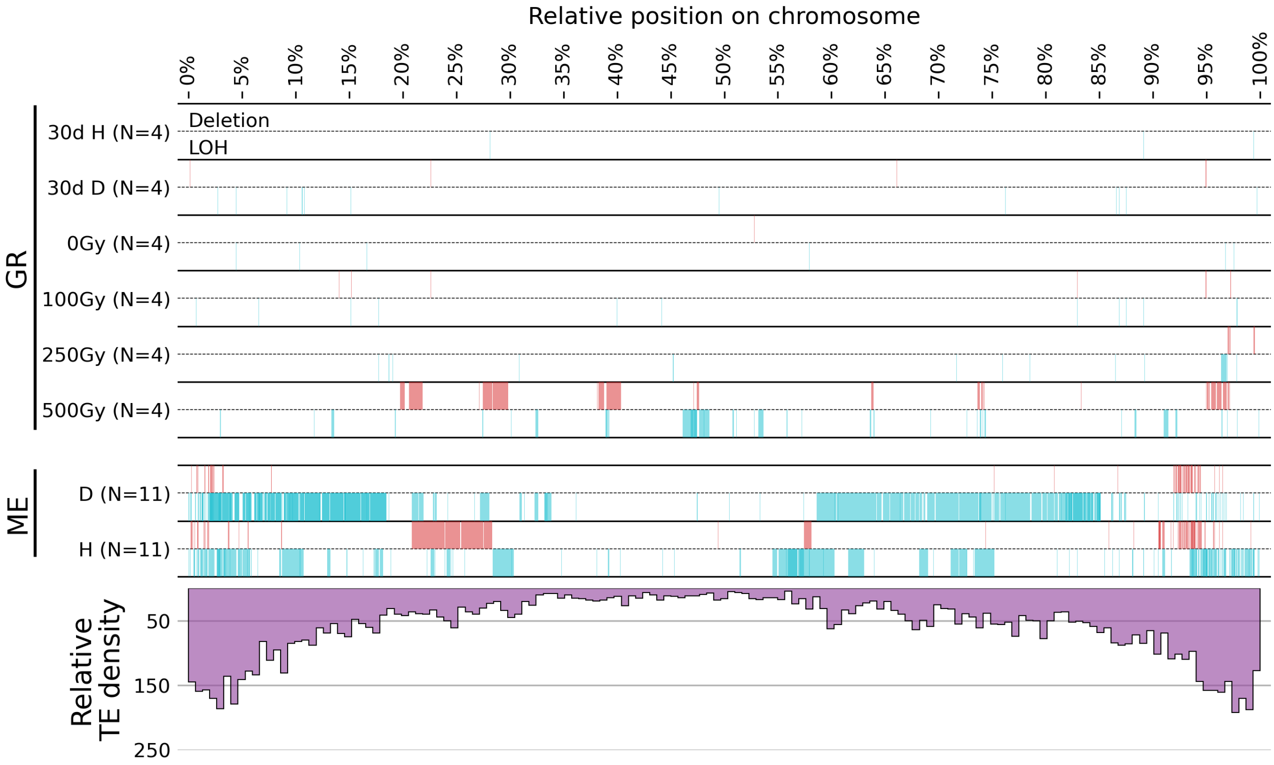


Supplementary Figure 9: Relative positions of LOH clusters categorized as either LOH (blue, no significant coverage drop compared to the ancestor) or deletions (red, categorized as deletion if $\Delta$ coverage $<$-20, indicating a coverage drop compared to the ancestor). Chromosomes are stacked on top of each other, with the position of each event represented proportionally to chromosome size.

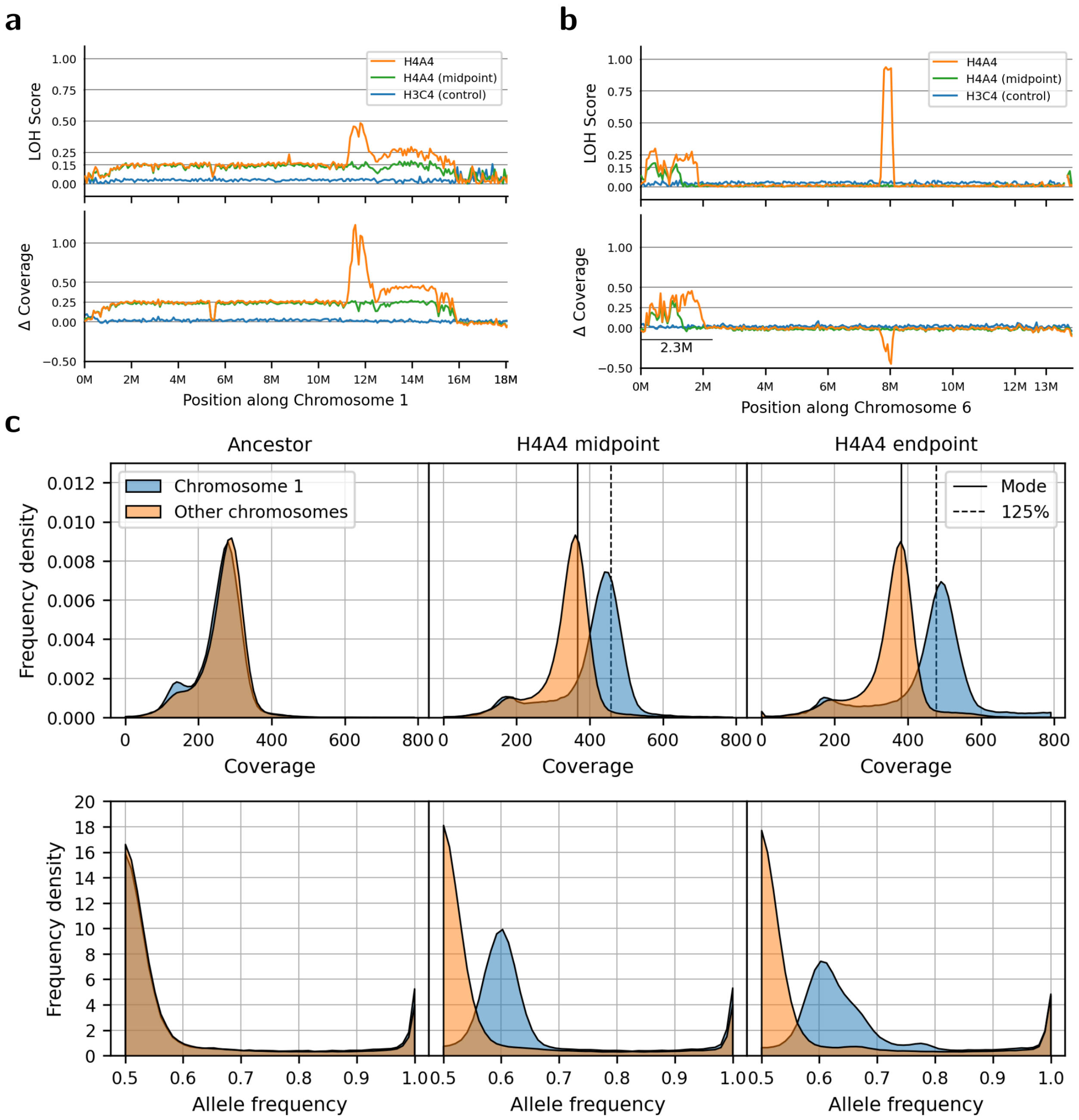


Supplementary Figure 10: **Whole genome duplication and additional chromosome 1 duplication in ME sample H4A4**. **a**: LOH score and Δ coverage along chromosome 1 in sample H4A4, showing a large duplication of the entire chromosome. One extremity appears unduplicated because the duplicated version of chromosome 1 is shorter. Additional copies have accumulated between generations 61 (midpoint) and 137 (endpoint) in the region between 11 and 16 Mb along the chromosome. **b**: LOH score and Δ coverage along chromosome 2, showing the duplication is limited to chromosome 1, and the observed shifts are not due to low sample coverage. **a-b**: ME sample H3C4 is shown as a control, in which no duplication was found on the same chromosomes. **c**: Comparison of coverage and allele frequency (AF) shifts between chromosome 1 and other chromosomes in the ancestor and H4A4 samples at both timepoints. The observed AF and coverage shifts are compatible with a WGD followed by an additional chromosome 1 duplication, with an allele ratio of $3/5 = 0.6$ (LOH score $2 \times |0.6 - 0.5| = 0.2$) and a coverage ratio of $5/4 = 1.25$ (Δ coverage $\sim$ +0.25). The LOH score is slightly lower than the theoretical value of $0.2$ because AF is based on the most frequent allele, which averages slightly above 0.5.

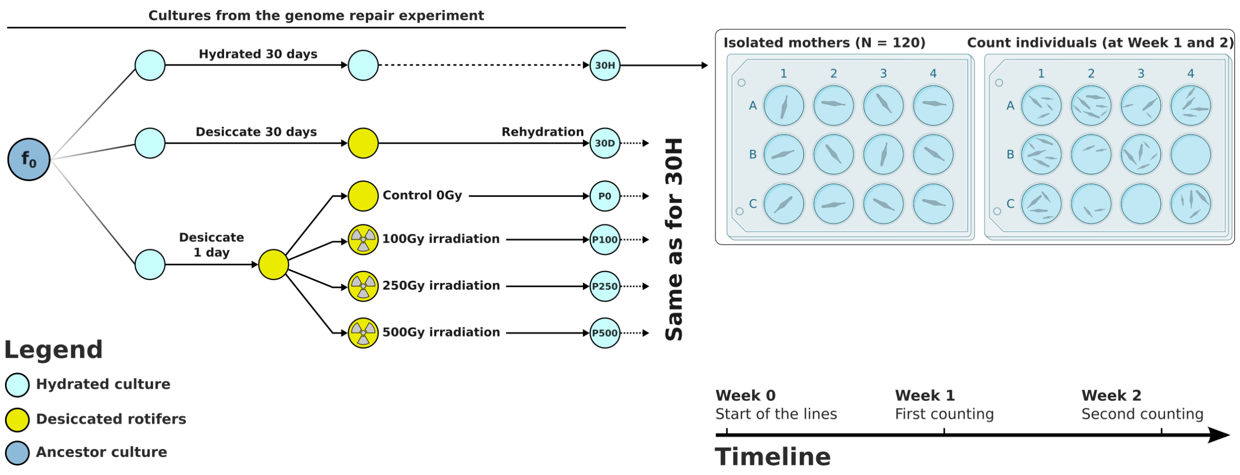


Supplementary Figure 11: Summary of the fertility experiment design. Post-treatment GR cultures were used to isolate 120 mothers. The number of active individuals from each mother was counted after 1 and 2 weeks under hydrated conditions.

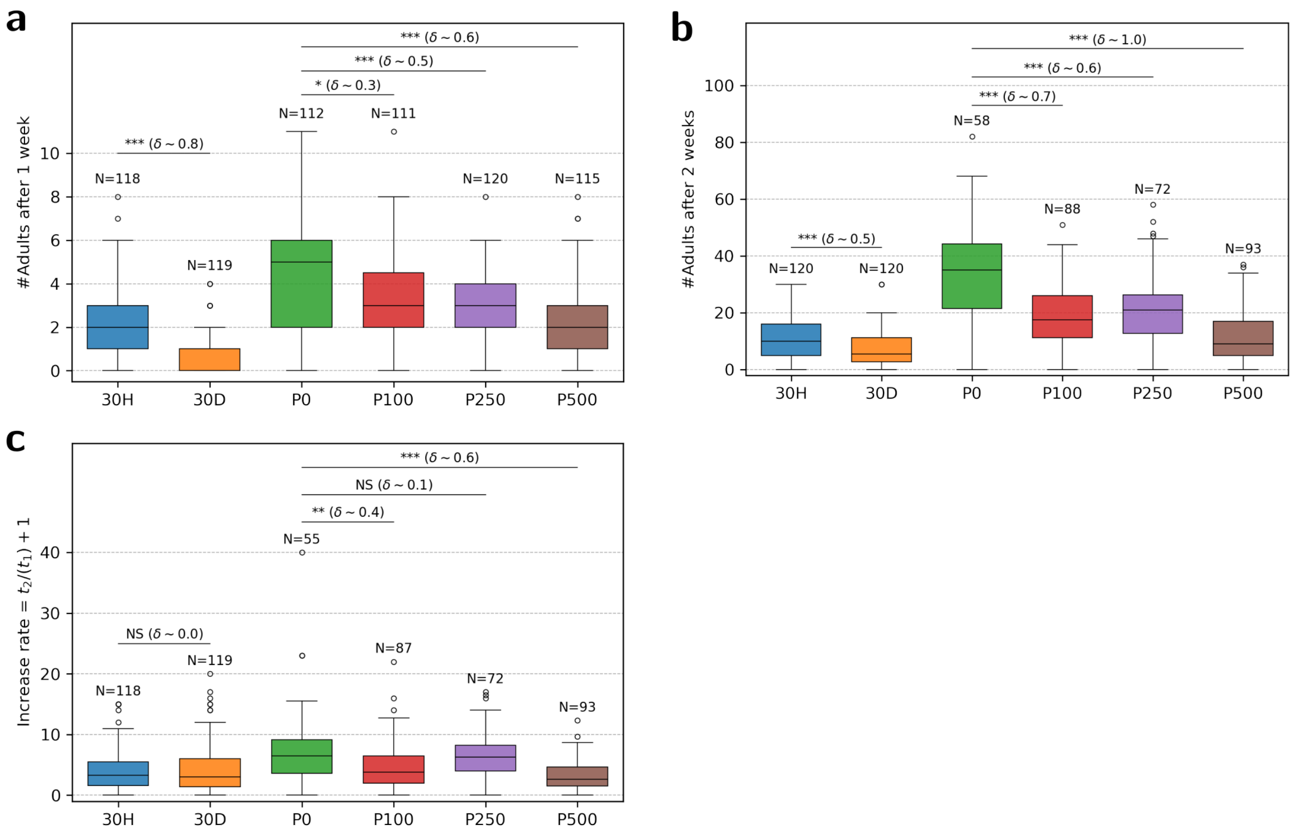


Supplementary Figure 12: **Effect of different conditions on fertility**. **a**: Number of adults counted in wells after week 1 ($t_1$). **b**: Number of adults counted in wells after week 2 ($t_2$). **c**: growth rate of adult numbers between week 1 and week 2. $N$: sample size, $\delta$: Glass' delta effect size statistics. NS: Non-significant; $* = p \leq 0.05$; $** = p \leq 0.01$; $*** = p \leq 0.001$. 30H: one month in hydrated culture, 30D: one month in desiccation, P0-500: proton irradiation dose received by cultures after one day in desiccation.

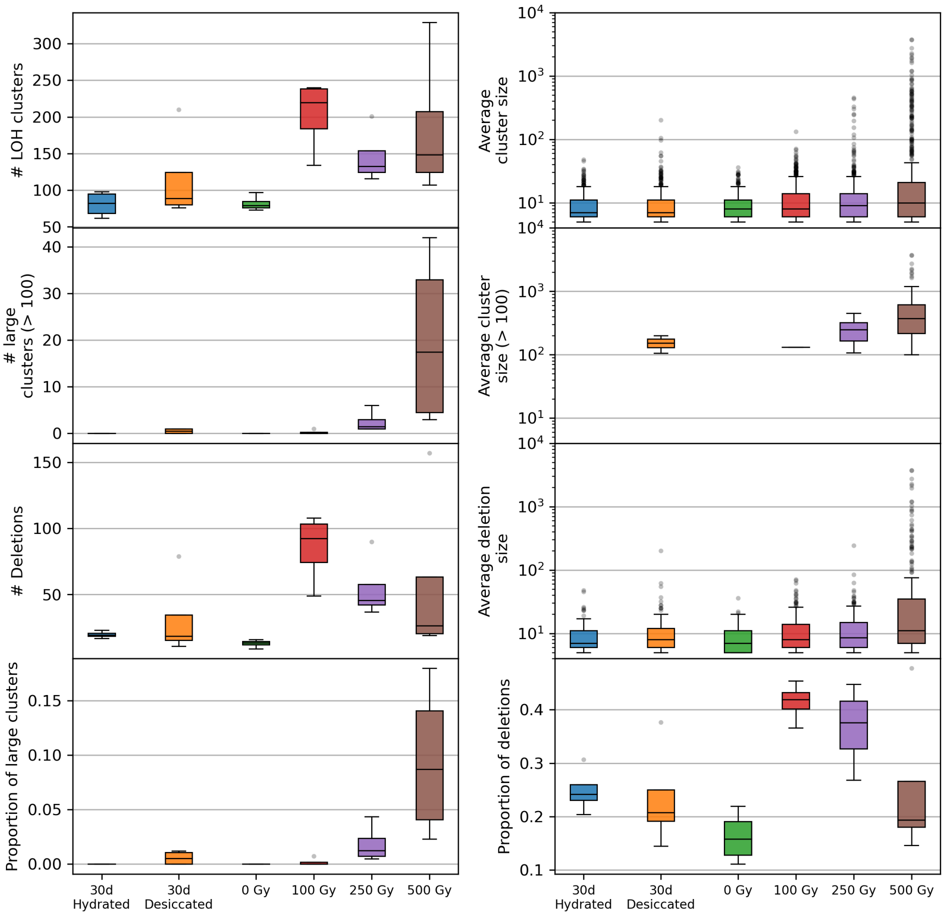


Supplementary Figure 13: Box plots showing the distribution of number and average size (measured by the number of ancestral heterozygous sites lost) under GR experimental conditions. The plots are organized as follows: the top shows clusters larger than 5 LOH sites, the middle displays clusters exceeding 100 LOH sites, and underneath shows the deletions ($\Delta$ coverage $<$-20). The bottom row, on the left, includes the distribution of the proportion of large clusters relative to all clusters, and on the right, the proportion of deletions to all clusters.

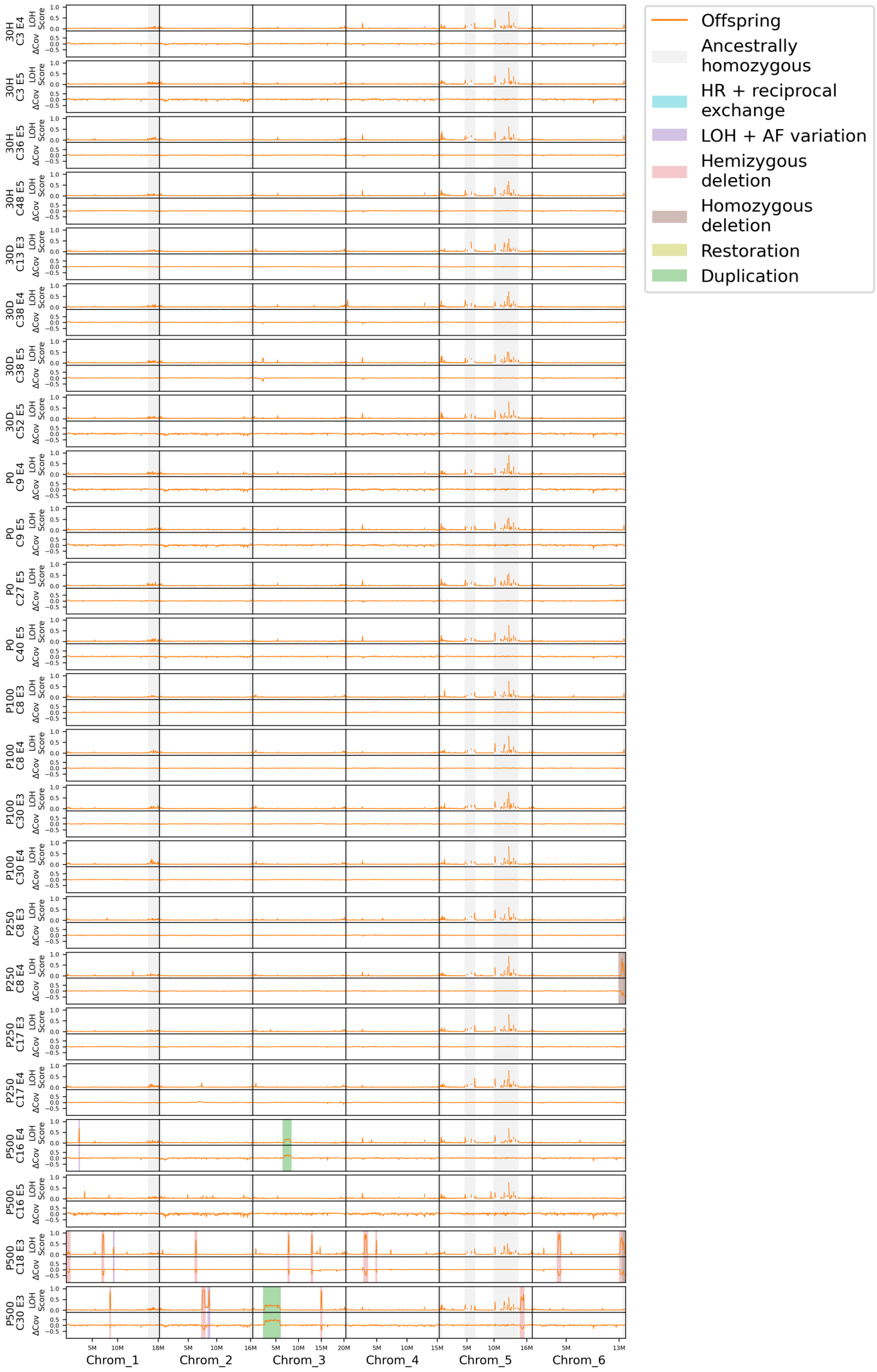


Supplementary Figure 14: LOH score and Δ coverage are displayed across all chromosomes for the GR samples. Large events (>50 Kb) are highlighted in color, blue for LOH, light red for hemizygous deletions, dark red for homozygous deletions, yellow for restorations and green for duplications. Grey areas represent regions that are ancestrally homozygous. Values are averaged over 250 non-overlapping windows along each chromosome, with window sizes comprised between 55.6 Kb (Chromosome 6) to 81.4 Kb (Chromosome 3).

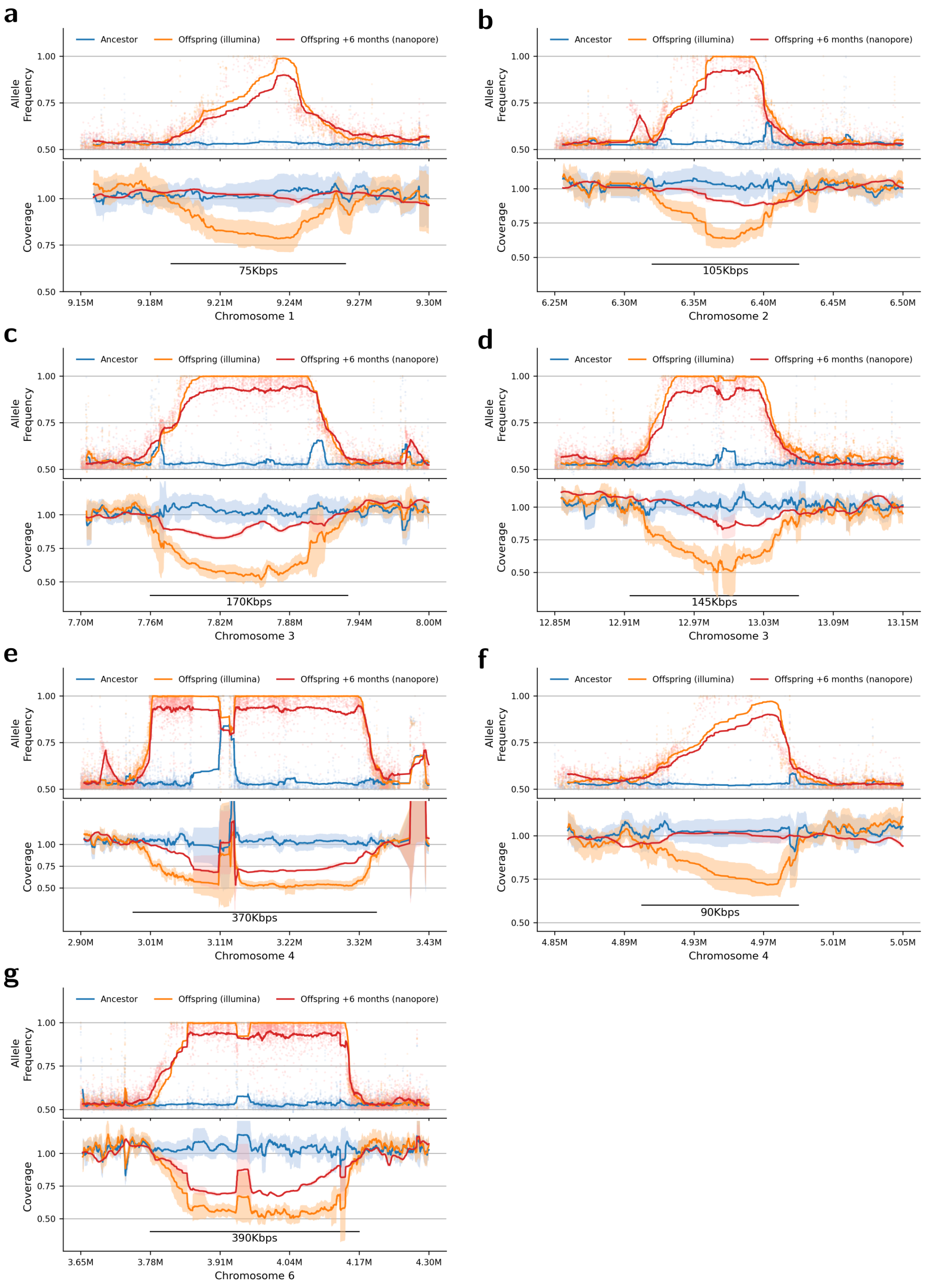


Supplementary Figure 15: **a-g**: The upper graph displays allele frequency (AF), while the lower graph illustrates coverage in regions where deletions were found in the GR 500 Gy sample P500 C18 E3. Both sequencing timepoints are compared to the ancestral Illumina sequencing (blue), with first an Illumina sequencing (orange) and then a Nanopore sequencing after 6 months of culture (red). The dots in the upper graph represent the AF of individual heterozygous sites, the line shows a sliding window average. The filled area of the lower graph corresponds to $\pm 1$ standard deviation around the average coverage in a sliding window.

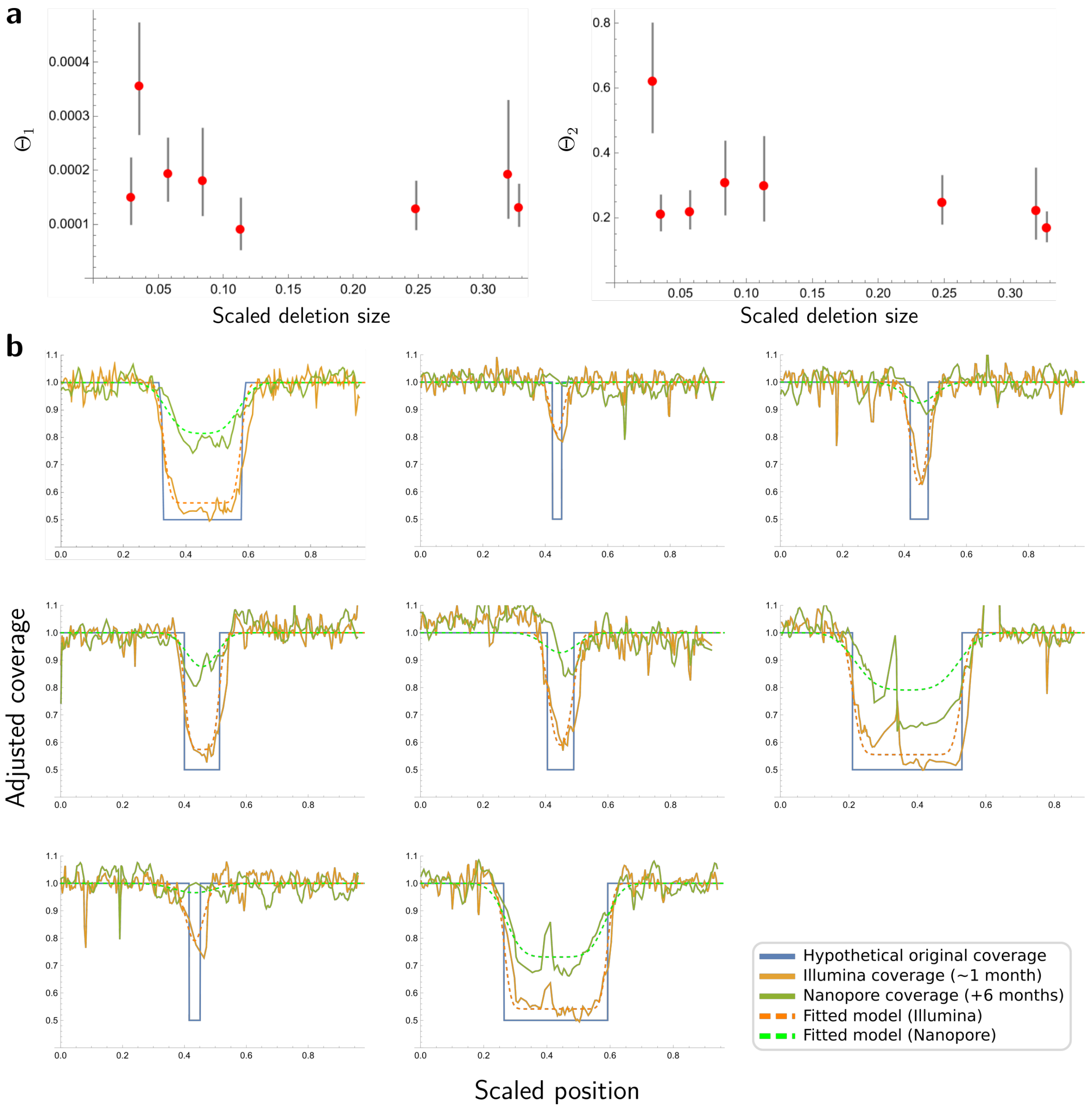


Supplementary Figure 16: **a**: Fitted values for parameters $\theta_1$ (degradation rate) and $\theta_2$ (repair rate) are represented, sorted by deletion size. These values are obtained by fitting the parameters from the diffusion model to the coverage data obtained from the GR 500 Gy sample at two time points: one month of culture (Illumina, orange) and six months of culture (Nanopore, green). **b**: The predicted coverage ($y-$axis, dashed lines) by position along the chromosome ($x-$axis) based on the fitted diffusion model for all deletions compared to the observed data (full lines) at both timepoints. The theoretical original deletion, where allele frequency reaches 1.0, is also shown (full blue line).

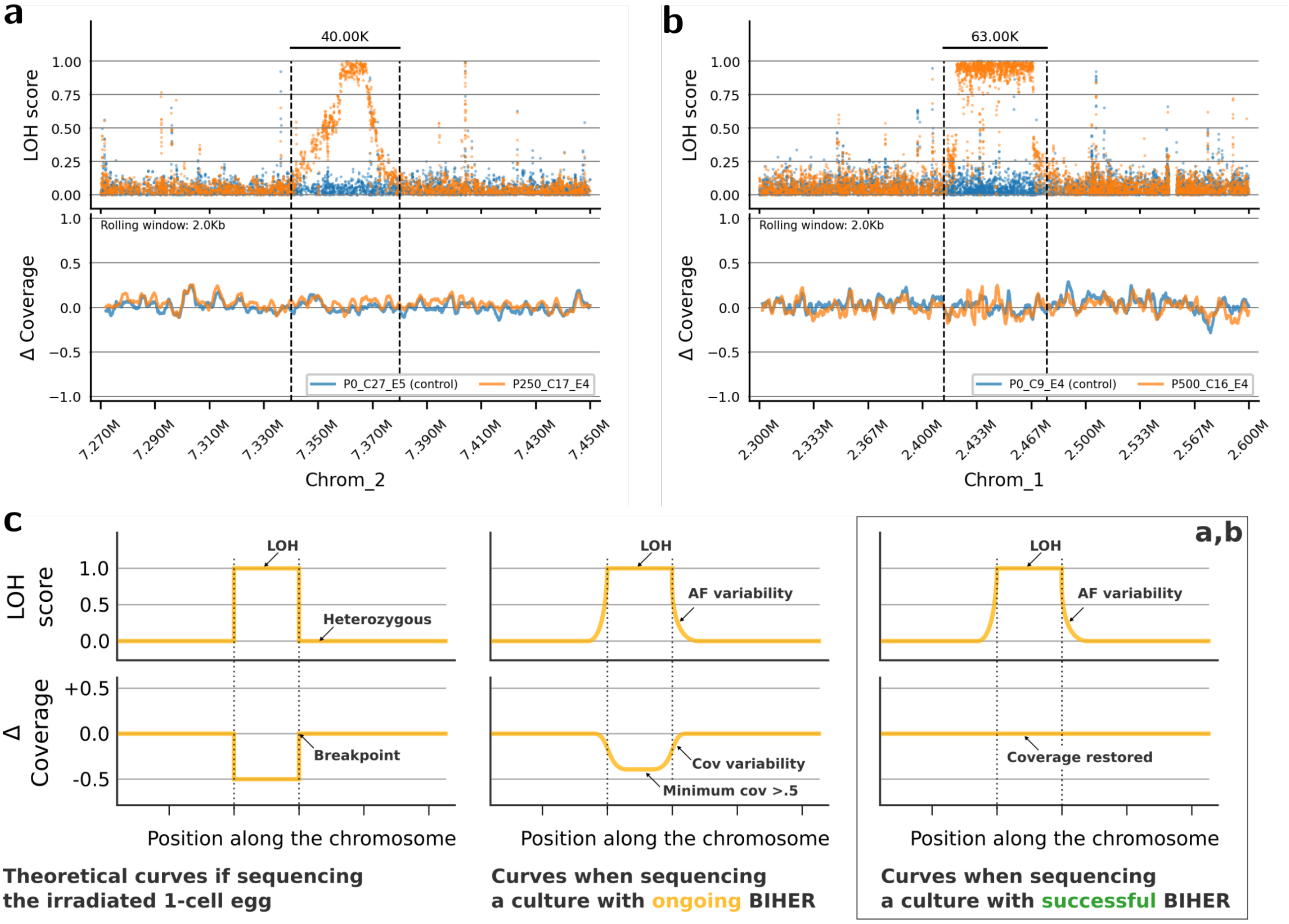


Supplementary Figure 17: **LOH tracts resulting from successful BIHER repair of deletions after GR 250 and 500 Gy IR**. **a-b**: The upper graphs show LOH scores, while the lower graphs display $\Delta$ coverage. **a**: LOH region identified on chromosome 2 in a 250 Gy IR sample, P250 C17 E4. **b**: LOH region identified on chromosome 1 in a 500 Gy IR sample, P500 C16 E4. **c**: Expected patterns of LOH score and $\Delta$ coverage at different stages of deletion repair by BIHER. Immediately after egg isolation, a fragmented chromosome deletion is present (top panel). During the culture period, ongoing BIHER and degradation create variability in fragment size among the cultured individuals. Over time, as more individuals in the culture fully repair the deletion, coverage increases (middle panel). In the two samples presented in panels **a** and **b**, at the time of sequencing, the LOH score still presents individual variability while coverage has been fully restored (bottom panel).

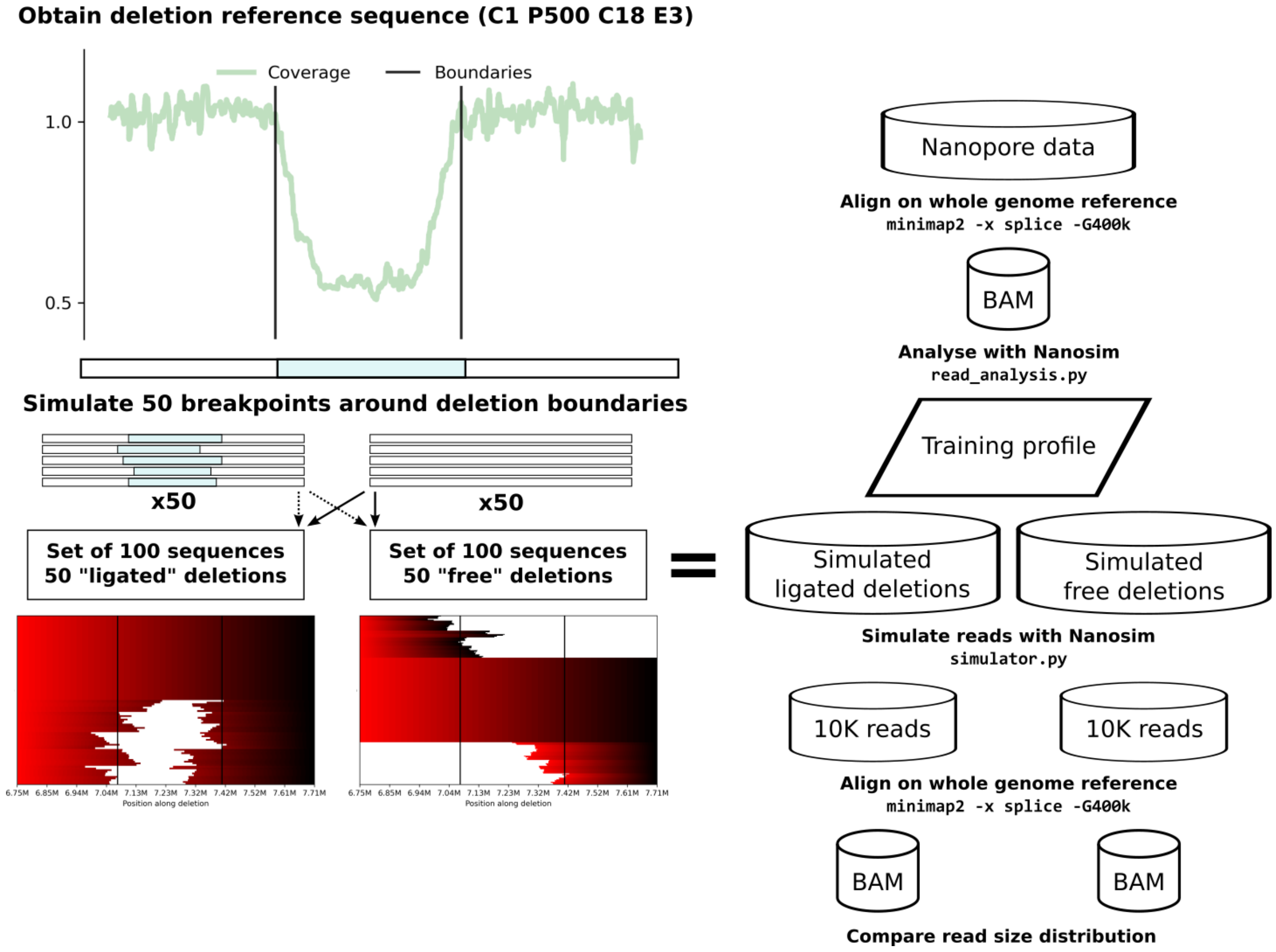


Supplementary Figure 18: **Methodology for simulating Nanopore reads**. A reference deletion was chosen from chromosome 1 in GR sample P500 C18 E3, in which few repetitive sequences were observed and for which coverage in the Nanopore data was not yet restored. From this reference sequence, we generated two sets of 100 deleted sequences, either ligated *in silico* or left fragmented. The breakpoint positions were determined using normal distributions centered around the actual LOH tract of the observed deletion, mimicking the process of resection and repair. In both sets, 50 sequences contained a deletion and 50 sequences were identical to the reference sequence (undeleted) to represent the presence of an unbroken homologous chromosome. The read profile from the real Nanopore dataset was obtained using `nanosim` and used to simulate 10,000 Nanopore reads from both sets of deleted sequences (fragmented or ligated *in silico*). The simulated reads, along with the real Nanopore reads, were aligned to the reference using `minimap2` allowing for considerable gaps in the alignment.

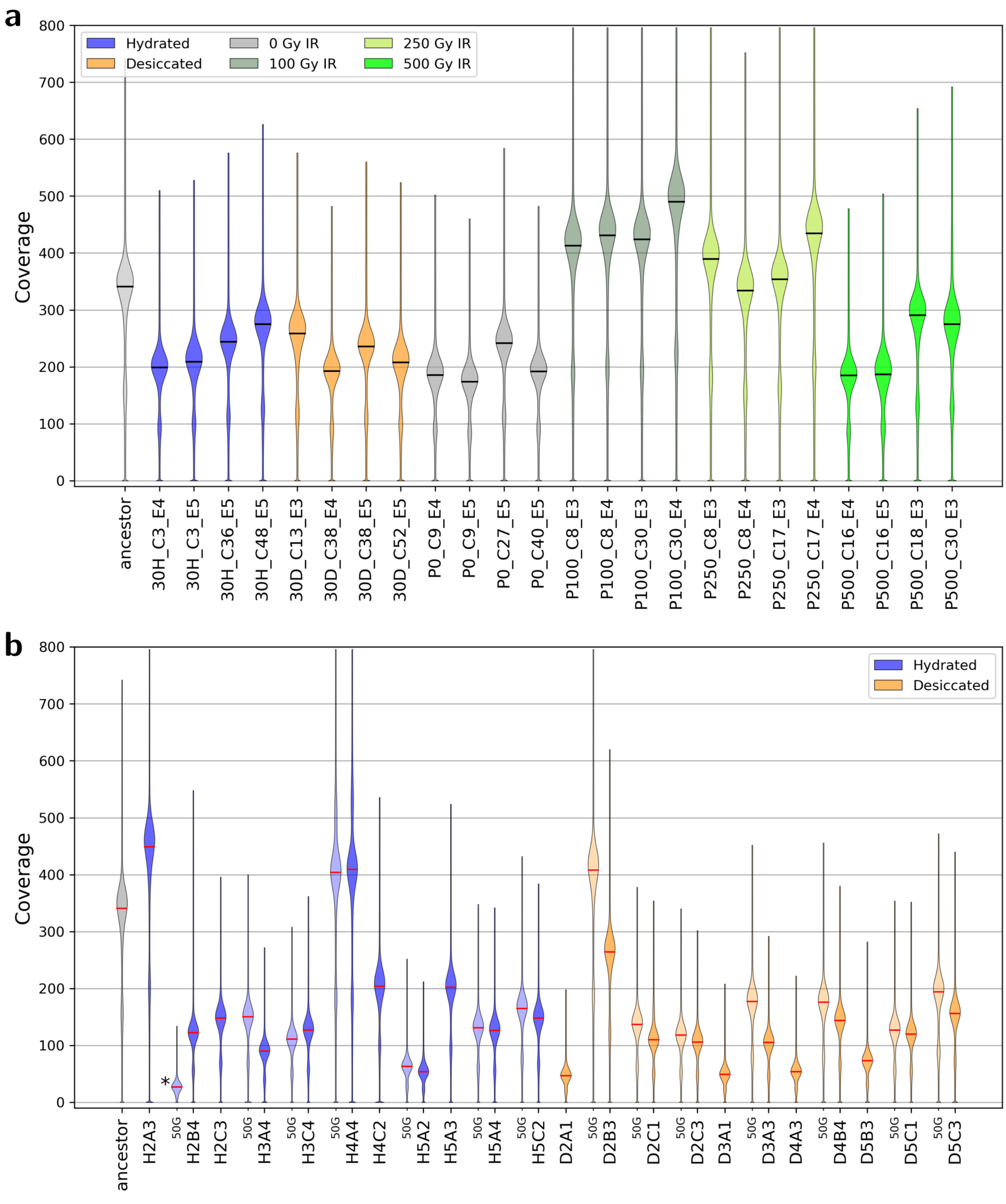


Supplementary Figure 19: **a-b** Coverage depth distribution per sample categorized per condition in color: blue for hydrated, orange for desiccated, gray for control 0 Gy IR, light green for 100 Gy IR, green for 250 Gy, dark green for 500 Gy. **a**: in the GR samples, average coverage ranges between 155x and 422x. **b**: in the ME samples, average coverage ranges between 26x and 391x. ME midpoints (50G) are shown in lighter shades and endpoints in darker shades. An asterisk (*) indicates the lowest coverage sample in ME (H2B4 midpoint) resulting from a technical problem during library sequencing.

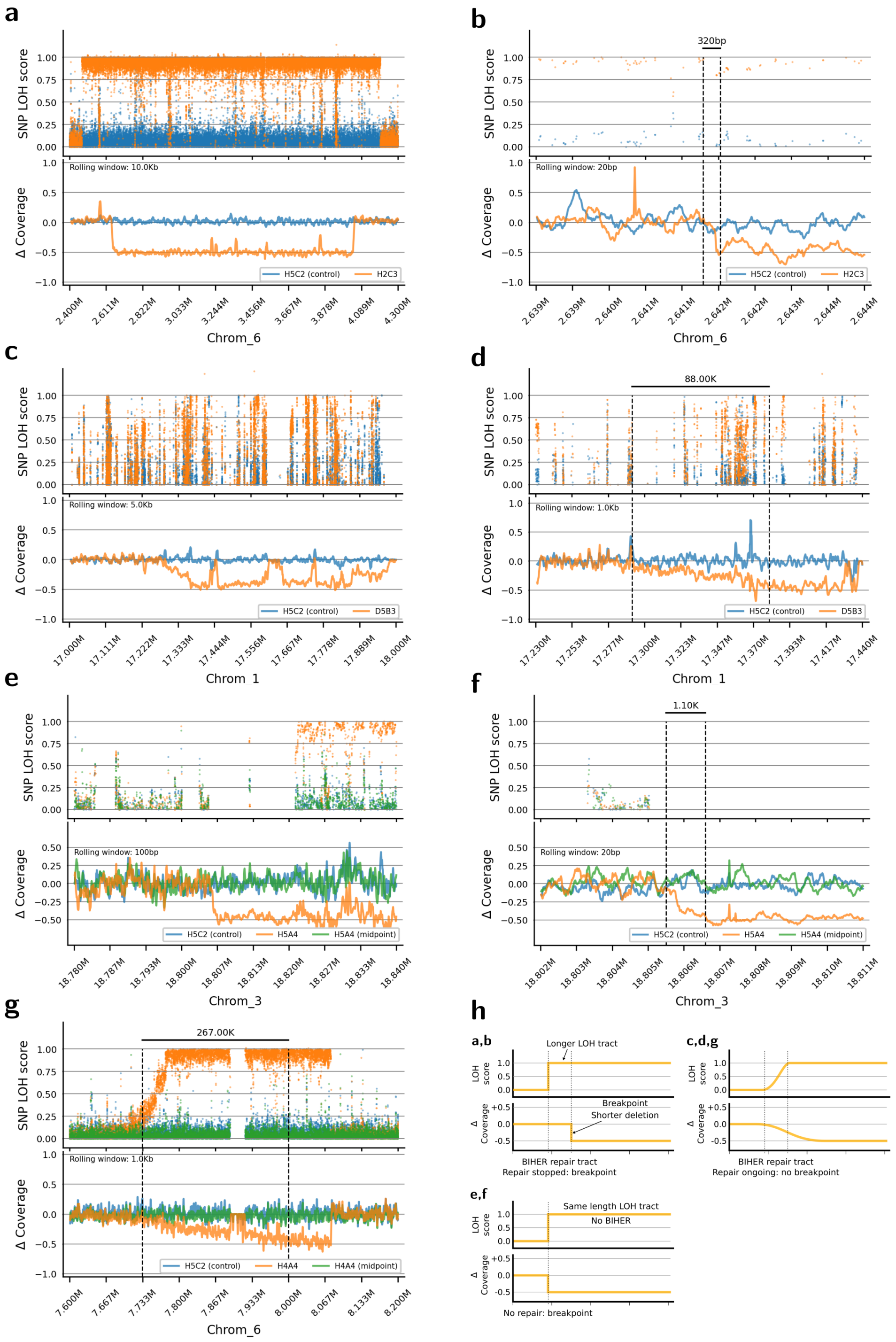

Supplementary Figure 20: Examples of coverage transitions observed in ME experiment samples. **a-g**: LOH score (upper graph) and $\Delta$ coverage (lower graph). **a**: Hemizygous deletion (1.4 Mb) on chromosome 6 of sample H2C3. The LOH tract extends past the coverage drop on both sides. **b**: Zoom on the left breakpoint of the deletion shown in panel **a**. **c** Homozygous deletion at the end of chromosome 1 of sample D5B3. **d**: Zoom on the left coverage transition (no breakpoints) of the deletion shown in panel **c**. **e**: Hemizygous deletion at the end of chromosome 3 of sample H5A4. **f**: Zoom on the deletion breakpoint of the deletion shown in panel **e**. **g**: Hemizygous deletion found on chromosome 6 of sample H4A4. The right side of the deletion has a breakpoint, and the left side of the deletion shows a progressive coverage transition. **h**: Representation of expected patterns of LOH score and $\Delta$ coverage in the samples, along with the corresponding repair hypothesis.

### Supplementary Tables

Table 1: Theoretical expectations for AF, coverage, LOH score and $\Delta$ coverage in cases of LOH due to RE or GC and hemizygous or homozygous deletions and duplications or restorations.

| | LOH | | Hemizygous Deletion | | Homozygous Deletion | | Restoration | | Duplication | |
|---|---|---|---|---|---|---|---|---|---|---|
| | $AB \rightarrow AA$ | | $AB \rightarrow A\emptyset$ | | $A\emptyset \rightarrow \emptyset\emptyset$ | | $A\emptyset \rightarrow AA$ | | $AB \rightarrow AAB$ | |
| | Ancestor **Anc** | Offspring **Off** | Anc | Off | Anc | Off | Anc | Off | Anc | Off |
| AF | 0.5 | 1.0 | 0.5 | 1.0 | 1.0 | 1.0 | 1.0 | 1.0 | 0.5 | 0.66 |
| Coverage | 1.0 | 1.0 | 1.0 | 0.5 | 0.5 | 0.0 | 0.5 | 1.0 | 1.0 | 1.5 |
| LOH score | 1.0 | | 1.0 | | 0.0 | | 0.0 | | 0.32 | |
| $\Delta_{Cov}$ | 0.0 | | -0.5 | | -0.5 | | +0.5 | | +0.5 | |
